# Integrated field and laboratory assessment of Swiss grapevine cultivar susceptibility to flavescence dorée reveals a central role for plant-vector interactions

**DOI:** 10.64898/2026.08.21.746194

**Authors:** Jasmine Cadena i Canals, Christophe Debonneville, Nathalie Dubuis, Isabelle Kellenberger, Michel Jeanrenaud, Olivier Viret, Stefano Bilotta, Alvaro Poretti, Guillaume Favre, Olivier Schumpp

## Abstract

Cultivar susceptibility strongly influences the epidemiology of vector-borne plant diseases, and understanding cultivar-specific variation can inform management strategies. This is particularly relevant for flavescence dorée, an incurable grapevine disease associated with a phytoplasma and transmitted by the leafhopper *Scaphoideus titanus*. In this study, we investigated the susceptibility of the main Swiss varieties, by combining controlled insect-mediated inoculation experiments with complementary field analyses conducted at progressively finer spatial scales. Together, these approaches allowed us to compare both infection probability and phytoplasma relative titre under standardised transmission conditions with disease incidence and relative titre under natural epidemiological conditions. For most cultivars, laboratory results were broadly consistent with field observations. However, a marked discrepancy emerged in the relative infection pattern between the two main grapevine cultivars grown in Switzerland: Chasselas and Pinot Noir. Under controlled conditions, they did not differ significantly in either their probability of infection or the phytoplasma relative titre, indicating no detectable difference in susceptibility to phytoplasma infection. In contrast, Pinot Noir consistently showed higher disease incidence than Chasselas under natural conditions. This pattern was observed across all spatial scales examined, from regional surveys to neighbouring vineyard plots, and was mirrored by higher phytoplasma relative titres. Importantly, under controlled conditions, *S. titanus* mortality during the one-week inoculation period was significantly higher on Chasselas than on Pinot Noir, indicating that Chasselas may provide a less favourable host for *S. titanus*. Together, these findings support the hypothesis that differences in field disease incidence between these cultivars may arise from differences in vector performance rather than intrinsic susceptibility to phytoplasma infection. This highlights the importance of considering plant-vector interactions, alongside susceptibility to infection, when assessing cultivar-specific vulnerability to vector-borne plant diseases.

**Author summary:** Grapevines are threatened by many diseases, including flavescence dorée, a serious disease associated with a phytoplasma, a wall-less bacterium that is transmitted between grapevines by the leafhopper *Scaphoideus titanus*. Because infected grapevines cannot be cured, disease management relies on removing infected plants, repeated insecticide applications against the vector, and annual surveillance. Here, we compared the susceptibility of the main grapevine cultivars grown in Switzerland by combining controlled transmission experiments with complementary field analyses based on ten years of disease surveillance data. Although the two main Swiss cultivars, Pinot Noir and Chasselas, were equally likely to become infected under controlled laboratory conditions, Chasselas showed a much lower disease incidence in vineyards. During the transmission experiments, we also found that insect mortality during the one-week inoculation period was substantially higher on Chasselas than on Pinot Noir. These findings suggest that Chasselas may be a lower-quality host for *S. titanus*, potentially contributing to the differences in disease incidence observed in the field. Further research is needed to test this hypothesis, which could help identify grapevine cultivars that are less suitable hosts for the insect vector and therefore less susceptible to flavescence dorée, ultimately informing breeding programmes and more sustainable disease management strategies.

## 1. Introduction

The susceptibility of different cultivars to pathogens and pests can vary significantly, and identifying and understanding this variability is crucial for effective disease management. This information can be used in breeding programmes or for developing more sustainable control methods (1). This becomes particularly critical for incurable diseases such as flavescence dorée (FD), since cultivar choice can represent an important lever to limit the spread of the disease. FD, associated with the Grapevine flavescence dorée phytoplasma (FDp) and transmitted by the leafhopper *Scaphoideus titanus* (2), is among the most damaging grapevine diseases (3,4). As no cure is available and FDp is regulated as a quarantine pathogen in the European Union (EU, 2019/2072) (5), control is based on insecticides against the vector, vineyard monitoring, and compulsory infected vine removal. In some regions, planting propagation material which has been hot-water treated is mandatory (6–8), since this treatment eliminates FDp and guarantees healthy vines (9,10). Mandatory control measures can slow down disease spread, but impose considerable agronomic, economic, and environmental costs (3,11). Moreover, FD has become so widespread in several European regions that eradication is no longer feasible, motivating the EU to authorise containment strategies instead (EU 2022/1630) (12), an approach that is also reflected in recent recommendations from the International Organisation of Vine and Wine (13). In this context, identifying less susceptible cultivars can provide a valuable lever for sustainable management, whether by planting them in high-pressure areas, by using them in breeding programs for tolerance or resistance, or by exploiting them to develop novel control strategies.

However, because grapevine cultivars are often strongly associated with specific wine-growing regions, exploiting reduced susceptibility requires the characterization of locally important cultivars. To date, only a limited fraction of grapevine cultivars has been characterized for susceptibility to FD, with available knowledge largely concentrated on cultivars studied in Italy, while cultivars grown in Switzerland remain largely unexplored (14). Cultivar differences in susceptibility can be expressed through variation in disease incidence, phytoplasma load, symptom expression or even plant recovery, a phenomenon whereby vines that showed FD symptoms and tested positive one year become spontaneously symptomless and FDp-negative in later years (15–17). Several field studies have reported cultivar-dependent variation in the aforementioned parameters (18–23), providing valuable insights under real viticultural conditions. Nevertheless, such studies are intrinsically context-dependent and influenced by uncontrolled factors that may bias results. These include vector pressure, which is strongly influenced by environmental context (24–26); temperature, which affects phytoplasma multiplication kinetics (27); and survey timing, as both phytoplasma titre and symptom expression differ throughout the season (22). Moreover, most studies have been conducted in single vineyards or geographically restricted areas rather than across broader FD-affected regions. As a result, they may not capture the full range of environmental conditions, vector pressures and climatic contexts that shape disease epidemiology, potentially confounding cultivar susceptibility with local environmental effects.

Laboratory experiments provide better standardization of environmental conditions and finer control over key factors such as pathogen exposure. They also enable the evaluation of a wide range of cultivars under comparable conditions, whereas cultivar diversity is often limited in the field by vineyard composition and disease distribution. However, controlled studies on FD remain challenging because of the difficulties associated with phytoplasma cultivation and transmission (28). To address these constraints, Eveillard *et al.* (2016) developed a workflow based on *ex vitro* plants and infectious *S. titanus* to assess cultivar susceptibility under laboratory conditions. They further demonstrated consistency between laboratory and field results for Cabernet Sauvignon and Merlot. Using a similar approach, Ripamonti *et al.* (2020a) evaluated cultivars from the Italian Piedmont region under laboratory conditions and compared the results with semi-field experiments conducted on grafted plants under controlled conditions.

Despite the numerous field studies and the limited investigations conducted under controlled conditions, comprehensive assessments of cultivar susceptibility remain scarce. Moreover, although large-scale FD surveillance data have been used to investigate field- and landscape-level factors associated with disease risk (31), cultivar susceptibility has not been specifically addressed by combining controlled inoculation experiments with field observations across an extensive FD-affected region. Addressing this knowledge gap is particularly relevant in Switzerland, where FD has progressively spread north of the Alps since its first detection in 2015 (32). Here, we addressed this gap by assessing the susceptibility of the main grapevine cultivars grown in FD-affected regions of Switzerland using complementary controlled inoculation experiments and multi-scale analyses of vineyard surveillance data.

## 2. Materials and methods

### 2.1. Plants and insects

#### 2.1.1. Plant material

The grapevine plants used in this study were obtained from cuttings prepared from canes collected in Agroscope vineyards. The cultivars and clones were as follows: Cabernet Sauvignon (unknown clone), Chardonnay (RAC 17), Chasselas (RAC 6), Gamay (ENTAV 565), Gamaret (RAC 14), Merlot (RAC 19) and Pinot Noir (RAC 12). Plants were grown in 1.3 L pots, trained to a single shoot and staked for support. Phytosanitary treatments included wettable sulphur and potassium bicarbonate against powdery mildew.

Prior to transmission assays, grapevine plants were screened by ELISA for major grapevine viruses, including Grapevine leafroll-associated viruses (GLRaV-1, GLRaV-2, GLRaV-3 and GLRaV-4), Arabis mosaic virus (ArMV), and Grapevine fanleaf virus (GFLV). Positive plants were excluded from the experiment.

Plants were trimmed 2 weeks (2023) or 4 weeks (2025) before the assays to induce lateral shoots growth. The young leaves produced were used for insect-mediated inoculation.

#### 2.1.2. Insect rearing

The laboratory colony of *Euscelidius variegatus* was established from individuals provided by IPSP-CNR (Turin, Italy), originating from a colony collected in Piedmont (33). The insects were reared in BugDorm-4 4S3074 insect cages (32.5 x 32.5 x 77cm MegaView Science Co., Ltd., Taichung, Taiwan) with two oat plants in phytotrons at 23 °C, 65% RH, 16 h photoperiod. *E. variegatus* were used as intermediate laboratory vectors to transmit the FDp from infected Madagascar periwinkle (*Catharanthus roseus*) to broad bean (*Vicia faba*), as *S. titanus* does not feed on periwinkle.

In 2023 and 2025, two-year-old grapevine canes were collected during winter pruning from three FD- and insecticide-free vineyards in the canton of Vaud, previously monitored for *S. titanus* flight activity. Canes were stored in darkness at 4 °C, covered with a plastic sheet and sprayed weekly with water to prevent desiccation. For insect emergence, wood from the three vineyards was pooled and placed in BugDorm-4 4S4590 insect rearing cage (47.5 x 47.5 x 93 cm MegaView Science Co., Ltd., Taichung, Taiwan) in a greenhouse (22 ± 2 °C, 60% RH day / 40% RH night, 14 h photoperiod). Wood was sprayed with water three times per week and a broad bean plant was provided in each cage as a food source for the nymphs. Plants were changed regularly when required during rearing.

#### 2.1.3. Obtaining FD-infectious S. titanus

An FDp isolate (FD-D, Genbank CP097583) is routinely maintained in Madagascar periwinkle plants in Agroscope’s quarantine greenhouse. Fifth-instar nymphs and adults of *E. variegatus* were allowed a one-week acquisition period on infected periwinkles, then transferred to broad beans. FDp presence in broad beans was confirmed by qPCR 6 weeks later. The isolate was maintained by cycling *E. variegatus* between infected and healthy broad bean plants.

To obtain infectious *S. titanus*, 5th instar nymphs were placed in a cage with an infected broad bean for a one-week acquisition period. After that, *S. titanus* were allowed for a 3-week latency period on healthy broad bean plants before being used for the transmission assays.

### 2.2. Assessment of cultivar susceptibility under controlled conditions

Under controlled conditions, cultivar susceptibility was evaluated using two complementary indicators: infection probability following experimental transmission of FDp by *S. titanus*, and phytoplasma relative titre in infected plants, expressed relative to the low-susceptibility reference cultivar Merlot (29). Infection status was determined by qPCR and used to estimate infection probability. Phytoplasma relative titre was quantified by qPCR using the ΔΔCt method (34) and calculated as the mean relative titre of all FDp-positive tissue samples collected from each infected plant.

#### 2.2.1. Transmission assays

Transmission assays were conducted in 2023 and 2025 using a protocol adapted from Éveillard et al. (29). Five experimental replicates were performed for Pinot Noir and Chasselas, four for Cabernet Sauvignon, Gamay, and Gamaret, and three for Merlot and Chardonnay, with 7 to 10 plants per experimental replicate at the start of the experiment. Experimental session corresponded to each inoculation block in which a subset of cultivars was exposed simultaneously to the same batch of potentially infectious insects. Details of experimental replicates, experimental sessions and numbers of plants are available in Table Supplementary 1 in S1 text. Session S01 is not represented in the analysed dataset because the corresponding experimental replicate did not meet the minimum proportion of infectious insects required for inclusion. Pinot Noir, Chasselas, Gamay, Gamaret and Merlot were chosen for the tests because they are the most widely grown cultivars in the canton of Vaud. Chardonnay and Cabernet Sauvignon were used as controls of the experience, as they have already been proven to be mildly and extremely susceptible before (29).

Four potentially infectious *S. titanus* were confined in insect-proof sleeves on a lateral shoot leaf, which was marked with a plant tie. Insects were left in place for one week; survival was checked the two following days and plants without surviving insects were discarded. After one week, the insects were collected from the sleeves, recorded as alive or dead, and stored in Eppendorf tubes containing 70% ethanol at −20 °C until tested for infectivity. After sleeve removal, plants were kept in the quarantine greenhouse for 15 more weeks. Experimental replicates in which less than 75% of the insects resulted infectious were discarded, and plants (experimental units) in which no insects were infected were also discarded.

Each experimental replicate also included two mock-inoculated (negative control) plants. For this purpose, four (in 2023) and three (in 2025) *S. titanus* reared on non-infected broad bean plants were confined in a sleeve for one week and handled in the same way as the infectious insects.

#### 2.2.2. Plant sampling

Plants were sampled 16 weeks after the start of the transmission assays. Four different leaves were collected per plant whenever possible: the inoculated leaf (IL), one leaf between the 3^rd^ and 5^th^ upper leaves (proximal upper leaf, PUL), another between the 8^th^ and the 10^th^ upper leaves (distal upper leaf, DUL) and one between the 3^rd^ and 5^th^ lower leaves (basal leaf, BL) (S1 Fig).

#### 2.2.3. Flavescence dorée detection

Insect and plant total nucleic acid extraction was performed with each insect individually and with approximately 0.5 g of leaf mid-vein tissue for plants following the protocol of Debonneville *et al.* (35). The presence of FDp and Bois noir phytoplasmas was then assessed by triplex quantitative PCR (qPCR) using a method adapted from Pelletier *et al.* (36). Insects were considered infectious if quantification cycle (Cq) for the FD gene was lower than 30, while plants were considered infected if quantification cycle was lower than 38. This threshold was chosen because 38.08 was the minimal Cq observed in the negative control plants group (10 plants out of 136 had a Cq between 38.08 and 40.75, the rest showed no amplification curve).

### 2.3. Assessment of cultivar susceptibility in the vineyards

Under field conditions, cultivar differences were evaluated using disease incidence and phytoplasma relative titre. Disease incidence was expressed as the density of FD-positive grapevines per hectare (FD+ vines ha⁻¹) for each cultivar. Annual vineyard surveillance data collected between 2015 and 2025 were used for the cantons of Vaud and Valais. Only the three regions with the highest FD incidence were included in the study (Fig 1): Lavaux and Chablais in the canton of Vaud, and Chablais valaisan and Central Valais in the canton of Valais. Chablais forms a continuous viticultural area across the cantonal border. Phytoplasma relative titre was quantified in the two main Swiss grapevine cultivars, Chasselas and Pinot Noir, from samples collected through the annual FD surveillance programme. Relative titre was determined using the ΔΔCt method, with the *Vitis* sp. chloroplast tRNA-L-F spacer as the reference gene (36) and Merlot as the reference cultivar. Because Merlot was underrepresented in the study area, additional reference samples were collected from vineyards in the canton of Ticino.

**Fig 1.**
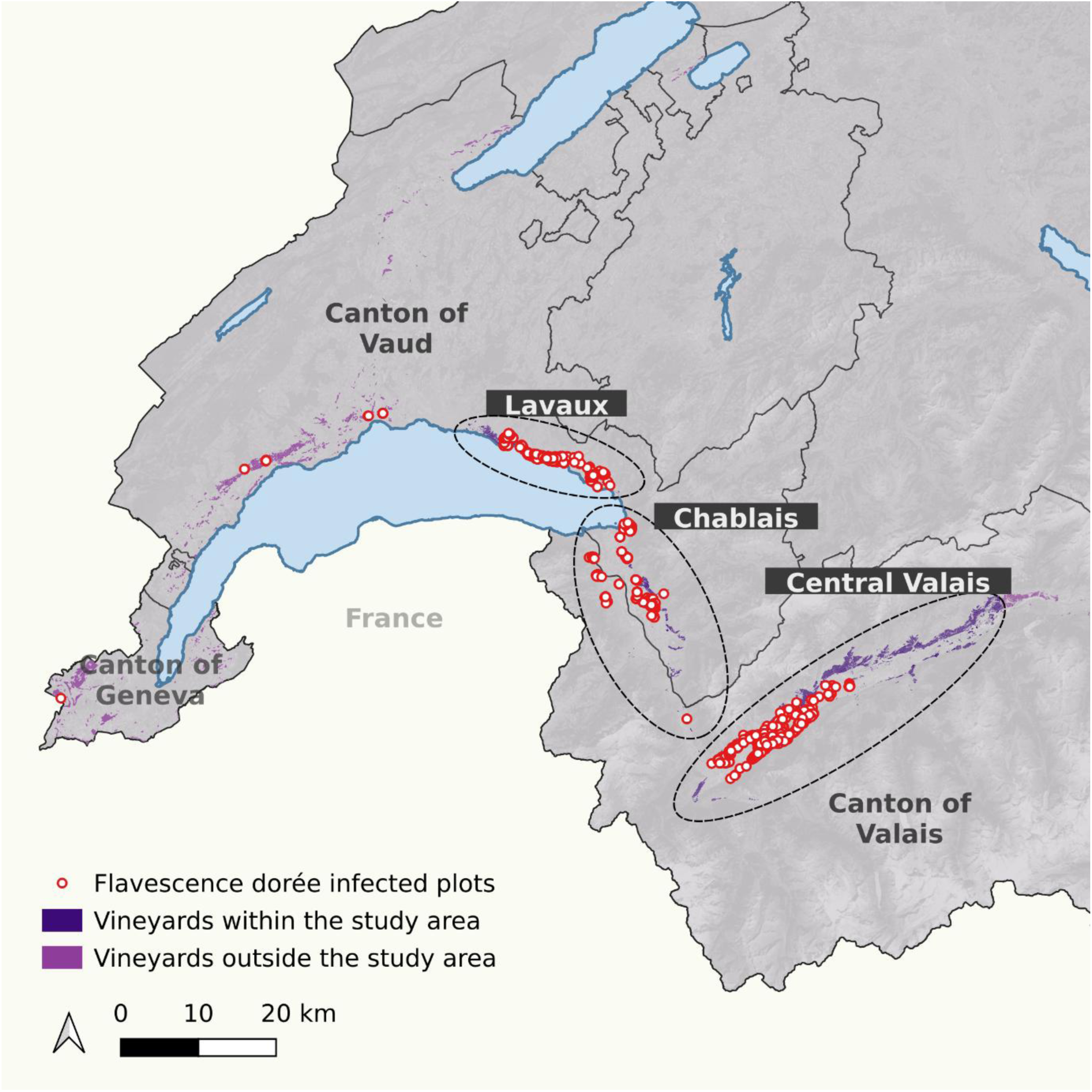
Study area and spatial distribution of Flavescence dorée-positive vineyards in western Switzerland. Dark purple polygons represent vineyards located within the study area, whereas light purple polygons indicate vineyards outside the study area. Red circles correspond to Flavescence dorée-positive plots recorded during annual surveillance between 2015 and 2025. Only vineyard areas in the cantons of Valais, Vaud, and Geneva are shown. The study focused on the viticultural regions with the highest Flavescence dorée prevalence, namely Lavaux, Chablais, and Central Valais (dashed ellipses).

To ensure that cultivar comparisons were not driven by spatial differences in disease pressure or cultivar distribution, analyses were conducted at three spatial scales (S2 Fig, S3 Fig and S4 Fig), each accounting for a different source of spatial heterogeneity: Macro: cultivar area in all three study regions (S2 Fig).

Meso: cultivar area within a 500 m buffer surrounding FD-positive grapevines. This distance corresponds to the perimeter defined in the Swiss federal directive (37) for establishing demarcated zones subject to mandatory insecticide treatments and therefore represents an at-risk area where both FDp and its insect vector are expected to occur (S3 Fig).

Micro: analysis restricted to Chasselas and Pinot Noir within 15 m of shared plot boundaries. Only buffer zones containing FD-positive grapevines were included. This design minimises spatial heterogeneity by comparing adjacent vineyard areas planted with different cultivars (S4 Fig).

The macro-scale analysis was conducted using surveillance data from both the cantons of Vaud and Valais. For the canton of Vaud, vineyard surface data and cultivar identities were obtained from the official 2022 vineyard cadastre, which provides georeferenced vineyard polygons and associated cultivar information. Georeferenced records of FD-positive grapevines collected during annual surveillance were also provided by the cantonal authorities. All spatial analyses were performed in QGIS 3.34.2-Prizren. The number of FD-positive grapevines associated with each cultivar was obtained by spatially joining FD-positive records to vineyard polygons, while cultivar surface areas were calculated directly from the cadastral polygons. For the canton of Valais, cultivar-specific vineyard surface areas and numbers of FD-positive grapevines were provided directly by the cantonal authorities and were therefore incorporated without additional spatial processing. Because many vineyard plots in Valais were planted with multiple cultivars, FD-positive grapevines recorded within these plots could not always be assigned unambiguously to a single cultivar. Consequently, all multicultivar plots were excluded from the Valais dataset, reducing the represented vineyard area from 4,372 to 2,961 ha and from 11,812 to 6,457 FD-positive grapevines.

The meso-scale analysis was conducted using only data from the canton of Vaud. For this analysis, 500 m buffer zones were generated around all FD-positive grapevines using the buffer tool in QGIS. Overlapping buffers were subsequently merged into a single at-risk area representing zones where both the disease and its vector were likely to occur. The resulting at-risk area was intersected with the 2022 vineyard cadastre to calculate cultivar-specific vineyard surfaces within the buffer zones. The number of FD-positive grapevines associated with each cultivar was obtained by counting FD-positive point records within vineyard polygons. These cultivar-specific FD-positive counts and vineyard surface areas were then used for subsequent analyses.

For the micro-scale analysis, only Chasselas and Pinot Noir in the canton of Vaud were considered. First, 2 m buffers were generated around vineyard polygons planted with either cultivar. Areas where these buffers overlapped were identified using the intersection tool and were considered contact zones between Chasselas and Pinot Noir vineyards. Subsequently, 15 m buffer zones were generated on both sides of these contact zones. Only buffer zones containing at least one FD-positive grapevine were retained. The vineyard cadastre was then intersected with the selected buffer zones to calculate cultivar-specific vineyard surfaces, and the corresponding numbers of FD-positive grapevines were extracted for statistical analyses.

To further minimise potential bias arising from vineyards that may have been planted with already infected grapevines, which could lead to FD-positive vines being overrepresented in particular cultivars, we examined a case study from a location where the origin of the outbreak was known (an infected Cabernet Dorsa vineyard). Vineyard planting densities were used to estimate the number of grapevines of each cultivar within the study area. FD incidence was then calculated as the proportion of infected grapevines relative to the estimated total number of grapevines of each cultivar, rather than to vineyard surface area.

### 2.4. Statistical analysis

All analyses were performed in R (version 4.5.2). Generalized linear models (GLMs) and generalized linear mixed models (GLMMs) were fitted using the glmmTMB package.

Model selection was restricted to a limited set of biologically motivated candidate models defined *a priori* according to the experimental design. Candidate error distributions and link functions were evaluated using Akaike’s Information Criterion (AIC) together with diagnostics based on simulated residuals generated with the *DHARMa* package, including assessments of uniformity, dispersion, zero inflation and outliers. Candidate fixed effects were then evaluated using likelihood ratio tests (LRTs) and AIC, whereas candidate random effects were evaluated using AIC together with the estimated random-effect variance. Estimated marginal means and contrasts were obtained using the emmeans package, using Tukey adjustment for all pairwise comparisons and Dunnett adjustment for planned contrasts against the reference cultivar Merlot, as appropriate. Statistical significance was assessed at α = 0.05. Complete analysis scripts, including model selection and diagnostic procedures, are available as described in the Data Availability statement.

To model the probability of plant infection in laboratory experiments, binomial models with logit, probit and complementary log-log (cloglog) links were evaluated. Because model fit was essentially identical among the candidate link functions (ΔAIC = 0.35), the complementary log-log link was retained owing to its closer correspondence with the underlying infection process, in which plant infection is expected to result from the accumulation of transmission opportunities provided by infectious insects. The number of infectious insects was included as an exposure offset to account for differences in infection pressure among plants. Experimental session, corresponding to each inoculation block in which a subset of cultivars was exposed simultaneously to the same batch of infectious insects, was evaluated as both a fixed and a random effect. Experimental session did not improve model fit when included as either a fixed or a random effect. The final model therefore included cultivar as the sole explanatory variable. Predicted infection probabilities were obtained from estimated marginal means at a standardised exposure of four infectious insects per plant and compared with the reference cultivar Merlot. An *a priori* targeted contrast between Pinot Noir and Chasselas was also performed.

Relative FDp titre in inoculated plants was analysed using generalized linear models. Gaussian, log-Gaussian and Gamma error distributions were evaluated. The number of infectious insects per plant, determined by qPCR testing of individual insects after the inoculation period, was included as an exposure offset to account for differences in inoculation pressure among plants. The final model consisted of a Gaussian model fitted to log-transformed relative titre values and included cultivar as the sole explanatory variable. Estimated relative titres were obtained from estimated marginal means at a standardised exposure of four infectious insects per plant. Comparisons with the reference cultivar Merlot and an *a priori* targeted contrast between Pinot Noir and Chasselas were then performed.

Insect mortality was analysed using a binomial GLMM. The model initially included cultivar, plant treatment (control = exposed to healthy insects, and inoculated = exposed to infected insects) and insect infection status as explanatory variables. Plant identity and experimental session were initially evaluated as random effects. Plant identity accounted for the non-independence of observations because healthy and infected insects from the same plant could contribute separate observations. Experimental session was subsequently retained as a fixed blocking factor, while plant identity was retained as a random intercept. Unsupported interactions and the insect infection status effect were removed during model selection. The final additive binomial GLMM therefore included cultivar, treatment and experimental session as fixed effects, and plant identity as a random effect. Estimated marginal means were used to obtain predicted mortality probabilities and associated 95% confidence intervals. Pairwise contrasts comparing each cultivar with the high-mortality reference cultivar Merlot were performed using Dunnett’s correction for multiple comparisons. In addition, an *a priori* targeted contrast was performed between Pinot Noir and Chasselas.

Data from the macro-, meso- and micro-scale vineyard analyses were analysed using negative binomial models (NB2) with a log link to account for overdispersion. Cultivar was included as an explanatory variable and vineyard area as an offset term so that estimates represent FD incidence per hectare. Estimated marginal means were computed and compared among cultivars using Tukey-adjusted pairwise comparisons. For the micro-scale analysis, a GLMM was fitted with cultivar as an explanatory variable, vineyard area as an offset term, and buffer zone identity as a random intercept.

The relative titre of field samples was analysed using a Gaussian linear model with identity link fitted to log-transformed data to account for the right-skewed distribution and heteroskedasticity of the data. The final model included cultivar as the sole explanatory variable. Estimated marginal means were computed, back-transformed to the original fold-change scale and pairwise comparisons were adjusted using Tukey’s method.

## 3. RESULTS

### 3.1. Cultivar susceptibility under controlled conditions

In 2023 and 2025, a total of 230 plants were exposed to four potentially infectious *S. titanus* for a week. After inoculation, the insects were tested for FDp infection status, and plants exposed only to non-infectious insects were excluded from the analysis. Across the entire experiment, the mean number of infected insects per plant was 3.24. Among the 230 exposed plants, 107 tested positive for FDp at the end of the experiment. The numbers of infected plants relative to the total number of exposed plants are shown in Fig 2.

**Fig 2.**
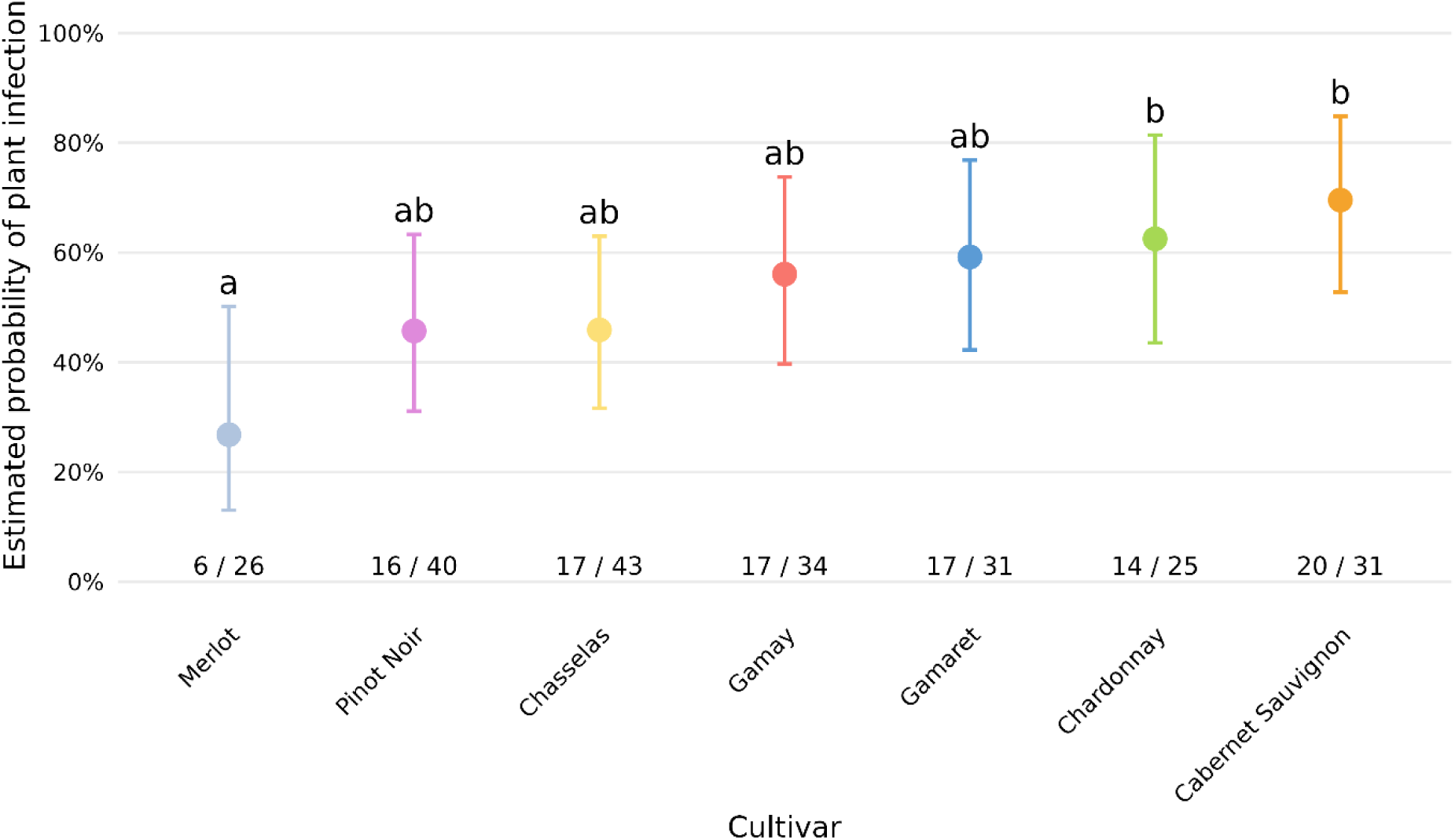
Model-estimated probability of FDp infection at 16 weeks after inoculation, standardised to an exposure of four infectious insects per plant. Points represent estimated probabilities (± 95% confidence intervals). Letters indicate significant differences from Merlot based on targeted Dunnett-adjusted contrasts (α = 0.05). Numbers indicate the number of infected grapevines out of the number of exposed grapevines per cultivar.

#### 3.1.1. Probability of infection of plants depending on cultivar under controlled conditions

The overall effect of cultivar on infection probability did not reach statistical significance (LRT χ² = 12.53, df = 6, p = 0.051). Contrasts comparing each cultivar with the low-susceptibility reference cultivar Merlot showed that Cabernet Sauvignon and Chardonnay were significantly more susceptible than Merlot, whereas the remaining cultivars formed an intermediate group (Fig 2, Tables Supplementary 2 in S1 Text). Among these cultivars, Gamay and Gamaret tended to show higher infection probabilities than Pinot Noir and Chasselas, but these differences were not statistically significant. The planned contrast between Pinot Noir and Chasselas was likewise not significant.

#### 3.1.2. Relative phytoplasma titre of infected plants under controlled conditions

Cultivar had no significant effect on phytoplasma relative titre (LRT: χ² = 9.73, df = 6, p = 0.137). Although model-estimated mean titres varied among cultivars, substantial within-cultivar variability resulted in wide confidence intervals with considerable overlap (Fig 3 and Tables Supplementary 3 in S1 text). Accordingly, Dunnett contrasts against the reference cultivar Merlot detected no significant differences among cultivars. Likewise, the planned contrast between Chasselas and Pinot Noir was not statistically significant (p = 0.311).

**Fig 3.**
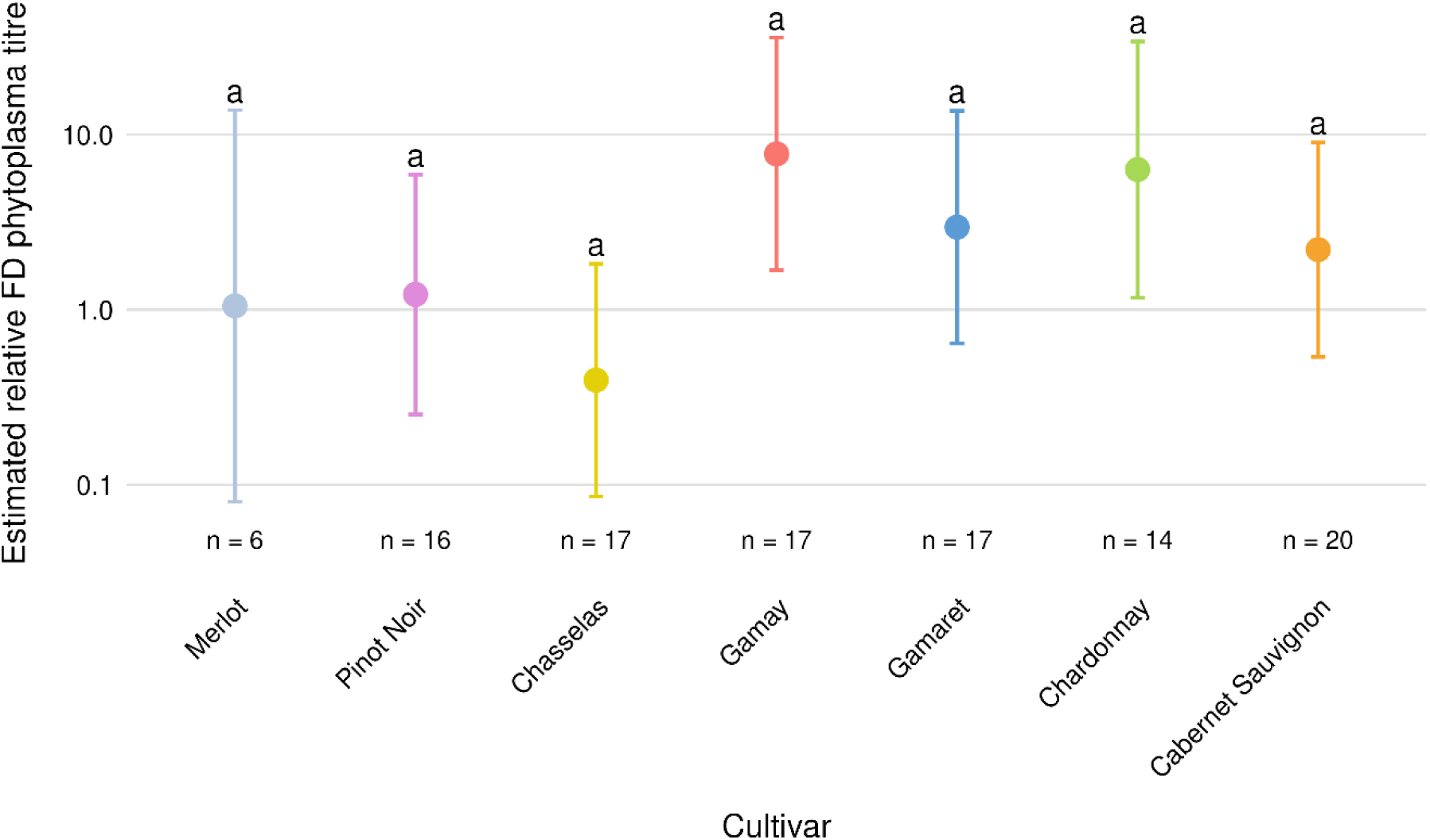
Estimated relative FDp titre in infected grapevines by cultivar following exposure to four infected insects per plant. Points represent estimated relative titres (± 95% confidence intervals). Estimates are expressed as ΔΔCt-derived fold changes relative to the reference cultivar Merlot. Letters indicate significant differences from Merlot based on targeted Dunnett-adjusted contrasts (α = 0.05). Numbers indicate the number of infected grapevines analysed per cultivar.

#### 3.1.3. Phytoplasma distribution in the plant

Phytoplasma distribution within infected plants was assessed by testing up to four leaves per plant (see Materials and Methods). A plant was considered infected when at least one sampled leaf tested positive for FDp. Among the 107 infected plants, the inoculated leaf tested positive in 70 cases, was negative in 21 cases, and was unavailable for analysis in 16 cases because it had abscised before sampling. These observations indicate that analysing multiple leaves increased the probability of detecting infection.

Sixteen weeks after inoculation, the inoculated leaf was the most frequently infected tissue across cultivars (Fig 4). In most cultivars, FDp was also commonly detected in proximal and distal upper leaves, indicating spread beyond the inoculation site, whereas basal leaves were infected much less frequently. Merlot differed markedly from the other cultivars, with no infected distal upper leaves and only one infected proximal upper leaf among infected plants, suggesting a much more restricted within-plant distribution of FDp.

**Fig 4.**
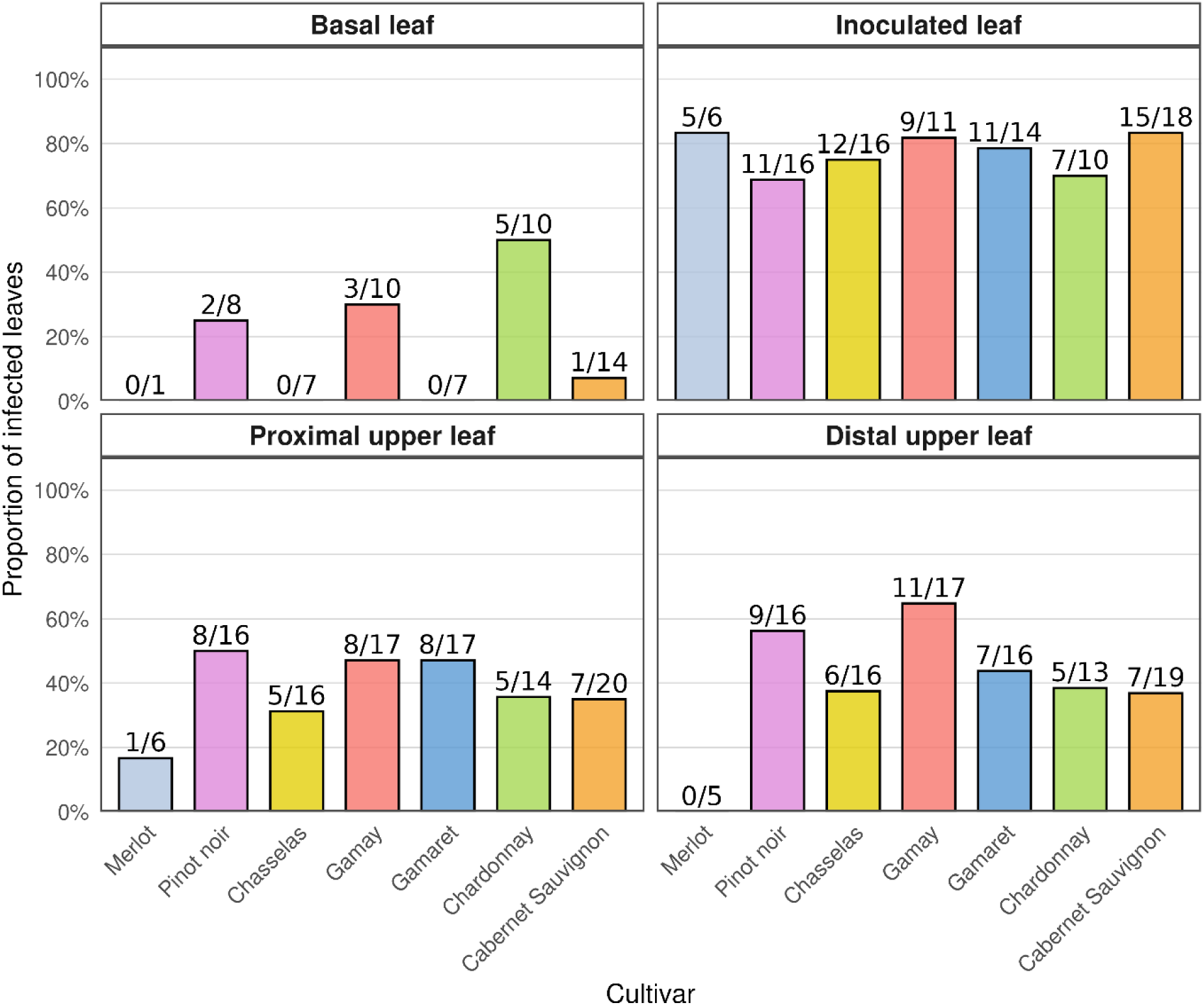
Distribution of FDp within positive grapevine plants across different leaf positions sampled 16 weeks after the start of the transmission assays for each cultivar. Bars represent the proportion of infected leaves, and labels above bars indicate the number of infected leaves relative to the total number of sampled leaves in positive plants. Leaf positions correspond to the basal leaf, inoculated leaf, proximal upper leaf, and distal upper leaf, as described in the Materials and Methods section.

#### 3.1.4. Insect mortality

Insect mortality differed significantly among cultivars (LRT χ² = 22.43, df = 6, p = 0.001). Mortality was also significantly higher in the inoculated treatment group (i.e., plants exposed to at least one infected insect) than in the control treatment group (i.e., plants exposed to non-infected insects) (LRT χ² = 14.34, df = 1, p < 0.001). In addition, experimental session had a significant effect on insect mortality (LRT χ² = 53.47, df = 8, p < 0.001), indicating substantial variation among inoculation blocks. Estimated mortality probabilities ranged from 35.7% (95% CI: 24.2-49.0%) in Cabernet Sauvignon to 75.5% (95% CI: 58.6-87.1%) in Merlot (Fig 5, Tables Supplementary 4 in S1 text). Compared with the high-mortality reference cultivar Merlot, mortality was significantly lower on Cabernet Sauvignon (Dunnett-adjusted p = 0.003), Chardonnay (p = 0.018), Gamay (p = 0.034) and Pinot Noir (p = 0.017), whereas Chasselas (p = 0.384) and Gamaret (p = 0.385) did not differ significantly from Merlot. The targeted comparison between the two principal grapevine cultivars grown in Switzerland showed that insect mortality was significantly lower on Pinot Noir than on Chasselas (odds ratio = 0.51, 95% CI: 0.28-0.91, p = 0.023).

**Fig 5.**
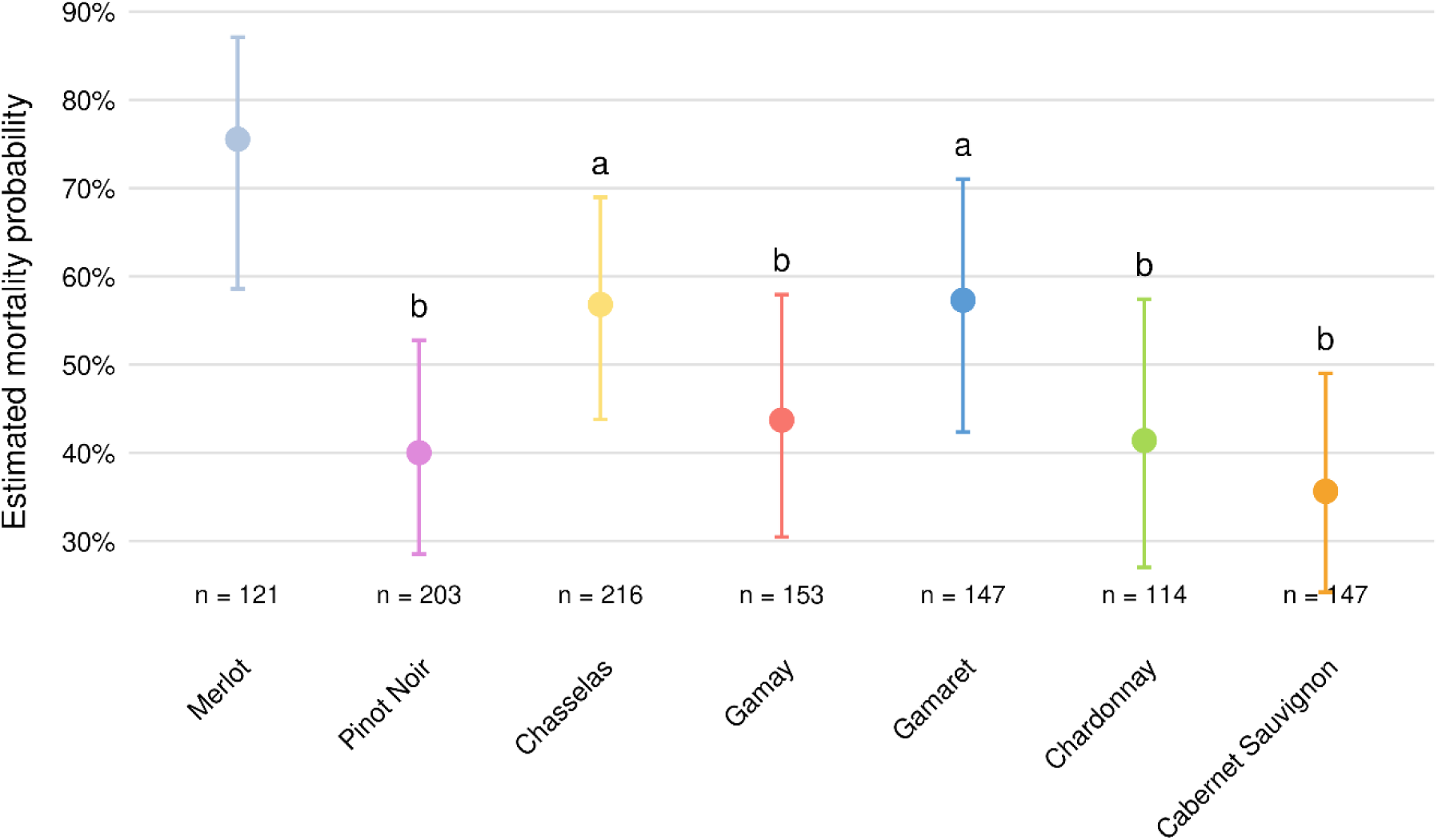
Estimated probability of insect mortality across grapevine cultivars. Points represent model-estimated mortality probabilities (± 95% confidence intervals). Different letters indicate cultivars that differ significantly from the reference cultivar Merlot based on Dunnett-adjusted contrasts (p < 0.05). Sample sizes (n = total number of insects) are shown for each cultivar.

### 3.2. Cultivar susceptibility in the vineyards

#### 3.2.1. Macro-scale study

The macro-scale analysis included 10,820 FD-positive grapevines distributed across 4,313 ha of monocultivar vineyards (69,213 plots). Analyses focused on the five most extensively cultivated cultivars in the study area: Chasselas, Pinot Noir, Gamay, Sylvaner and Arvine. Together, these cultivars accounted for 8,123 FD+ grapevines and represented 3,202 ha (49,418 plots).

Cultivar had a highly significant effect on the estimated density of FD-positive grapevines (LRT: χ² = 447.98, df = 4, p < 0.001). Gamay showed the highest estimated density (5.29 FD-positive plants ha⁻¹, 95% CI: 4.08-6.85), followed by Pinot Noir (2.56, 95% CI: 2.11-3.10), and the two cultivars differed significantly. Chasselas showed an intermediate estimated density (0.27, 95% CI: 0.22-0.33). Arvine had a lower estimated density (0.15, 95% CI: 0.07-0.28) and did not differ significantly from either Chasselas or Sylvaner. Sylvaner showed the lowest estimated density (0.03, 95% CI: 0.01-0.08). Tukey-adjusted pairwise comparisons identified four partially overlapping statistical groups among the five cultivars (Fig 6, Tables Supplementary 5 in S1 text).

**Fig 6.**
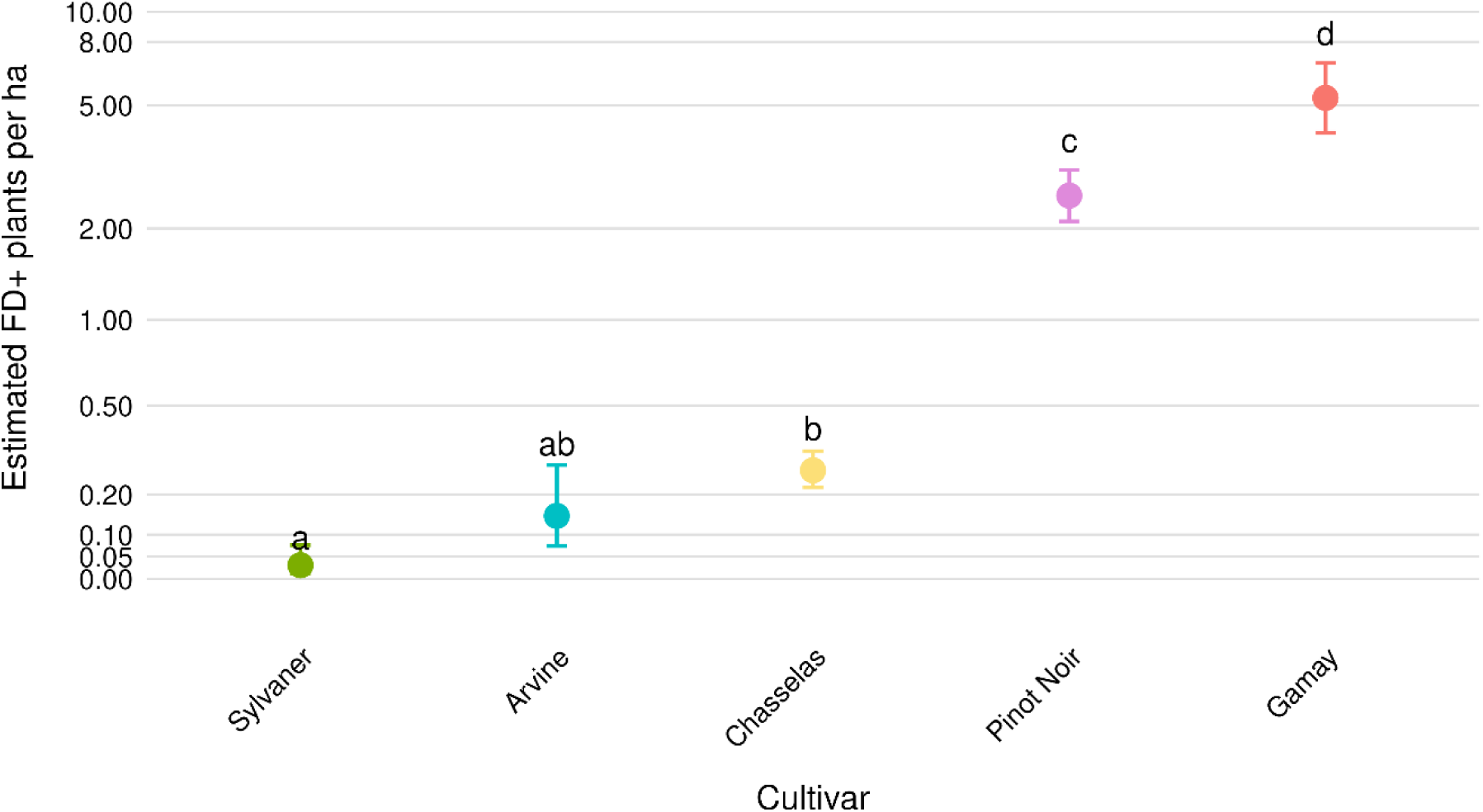
Estimated FD+ plants per hectare for the five main grapevine cultivars included in the macro study zone (Lavaux, Chablais and Central Valais). Points represent model-estimated marginal means and error bars indicate 95% confidence intervals. Different letters indicate significant differences among cultivars based on Tukey-adjusted pairwise comparisons (α = 0.05).

#### 3.2.2. Meso-scale study

The meso-scale analysis included 2,937 FD-positive grapevines distributed across 664 ha of vineyards within the 500 m at-risk zones in the Lavaux and Chablais regions of the canton of Vaud. Analyses focused on the five most extensively cultivated cultivars of the plots included in the meso-scale study: Chasselas, Pinot Noir, Gamay, Gamaret and Merlot. As with the macro-scale study, cultivar had a highly significant effect on the estimated incidence of FD-positive grapevines (LRT: χ² = 190.5, df = 4, p < 0.001). Compared to the macro-scale analysis, differences among represented cultivars were less pronounced, with Pinot Noir, Gamay and Gamaret belonging to the same statistical group, whereas Chasselas and Merlot showed significantly lower estimated incidences (Fig 7, Tables Supplementary 6 in S1 text). Because Chasselas still represented more than 70% of the vineyard surface within the at-risk zones, a finer-scale analysis was subsequently performed.

**Fig 7.**
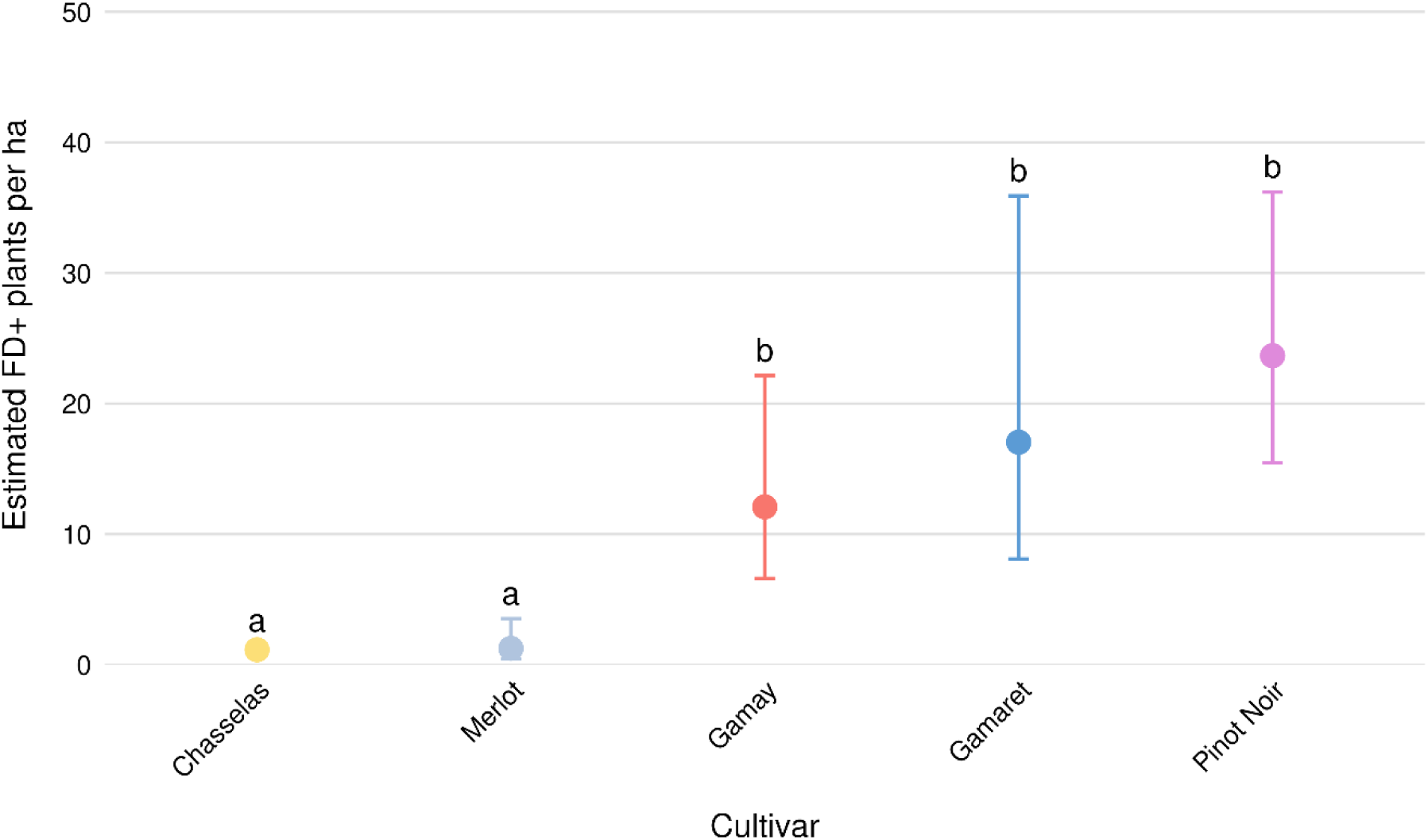
Estimated FD+ plants per hectare for the five main grapevine cultivars included in the 500 m at-risk study zone (Lavaux and Chablais in the canton of Vaud). Points represent model-estimated marginal means and error bars indicate 95% confidence intervals. Different letters indicate significant differences among cultivars based on Tukey-adjusted pairwise comparisons (α = 0.05).

#### 3.2.3. Micro-scale analysis

The micro-scale analysis included 65 buffer zones located along Chasselas-Pinot Noir plot boundaries in Lavaux and Chablais, canton of Vaud. These buffer zones encompassed 27.94 ha and contained 1,011 FD-positive grapevines across all grapevine cultivars. After restricting the dataset to Chasselas and Pinot Noir, the final analysis included 25.16 ha and 884 FD-positive grapevines. Strong spatial heterogeneity was detected among buffer zones, as including buffer zone as a random effect substantially improved model fit (AIC = 557.9 vs. 588.3). After accounting for this spatial heterogeneity, cultivar had a highly significant effect on the estimated incidence of FD-positive grapevines (LRT: χ² = 26.82, df = 1, p < 0.001), with Pinot Noir showing an estimated incidence approximately 6.6 times higher than Chasselas (28.27, 95% CI: 15.86-50.39 vs. 4.30, 95% CI: 2.26-8.19 FD-positive plants ha⁻¹; p < 0.001) (Fig 8, Tables Supplementary 7 in S1 text).

**Fig 8.**
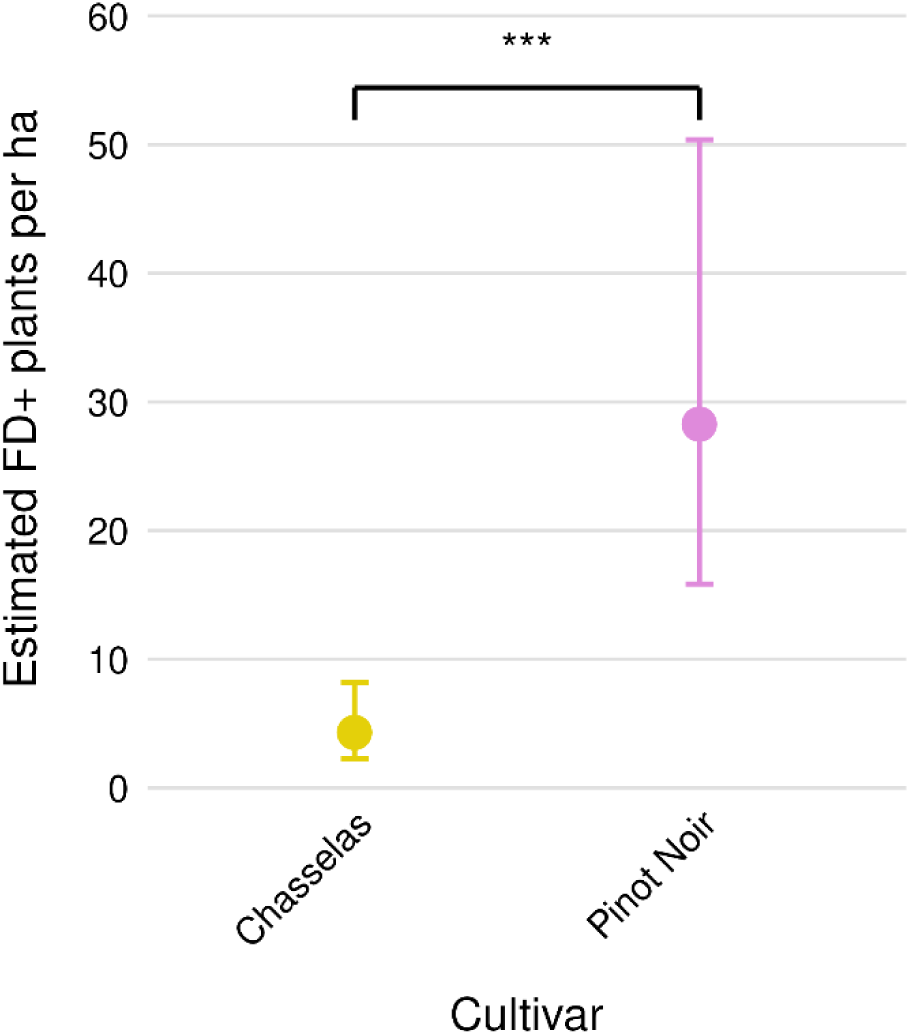
Estimated FD+ plants per hectare for the two main grapevine cultivars included in the micro-scale study (Lavaux and Chablais in the canton of Vaud). Points represent model-estimated marginal means and error bars indicate 95% confidence intervals. Asterisks indicate significant differences among cultivars based on Tukey-adjusted pairwise comparisons (α = 0.05).

#### 3.2.4. Case study

The case study was designed to illustrate the natural in-field incidence of FD in Chasselas and Pinot Noir in a context where the initial inoculum source was known not to be in one of these two cultivars, namely a nearby Cabernet Dorsa vineyard containing infected grapevines (Fig 9). Furthermore, the vine densities were known, enabling the FD+ cases to be reported against the theoretical number of at-risk vines. In Pinot Noir, 322 grapevines out of 6,020 (5.35%) were found positive, whereas only 31 out of 13,783 (0.22%) were found positive in Chasselas. Figure 9B shows that FD-positive grapevines accumulated predominantly within Pinot Noir vineyards, whereas comparatively few cases were detected in neighbouring Chasselas vineyards.

**Fig 9.**
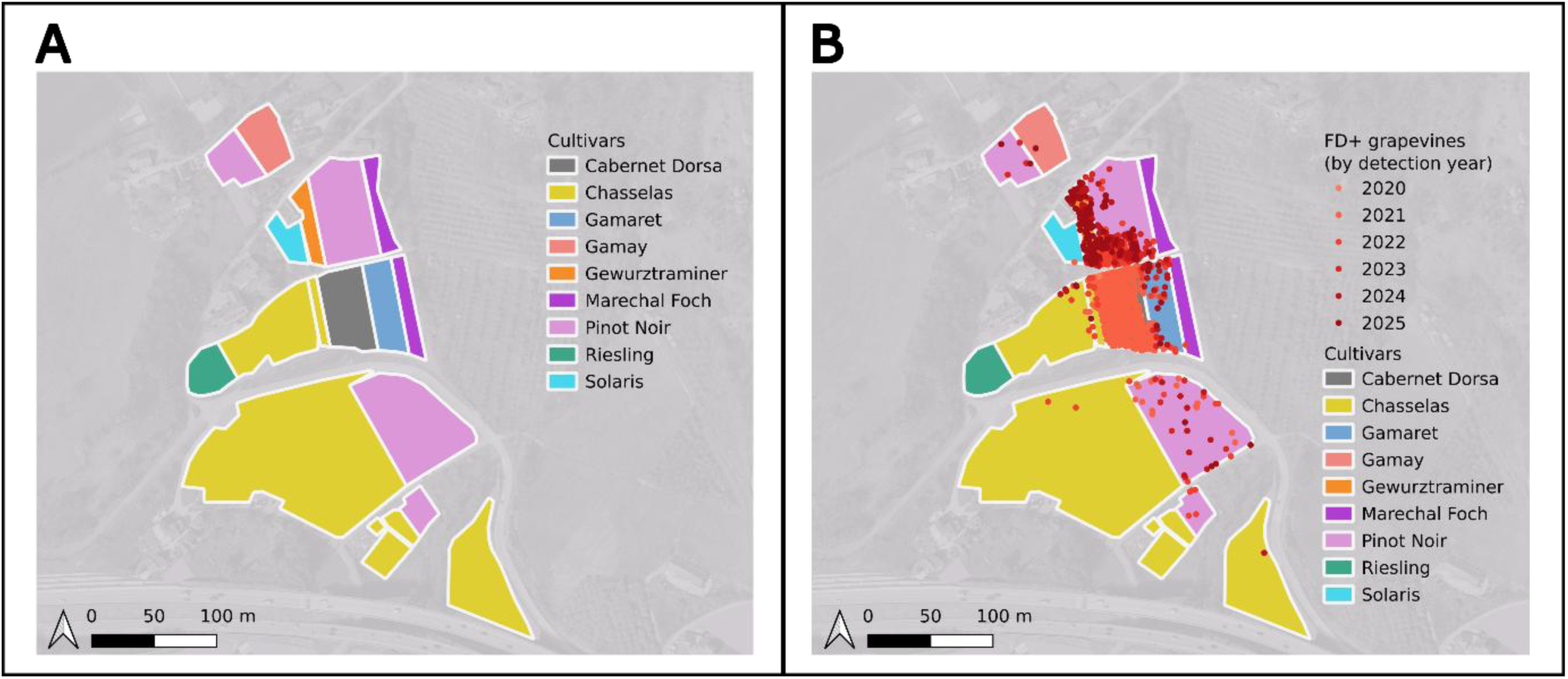
Case-study vineyard used for susceptibility assessment. (A) Spatial arrangement of the different grapevine cultivars. (B) Distribution of FD+ grapevines recorded from 2020 to 2025, coloured according to the year of detection.

#### 3.2.5. Relative FDp titre in the vineyards

The relative FDp titre was compared among field samples of Merlot (n = 59), Chasselas (n = 47) and Pinot Noir (n = 70). The LRT revealed a highly significant effect of cultivar (χ² = 100.39, df = 2, p < 0.001). Estimated marginal means and Tukey-adjusted pairwise comparisons showed significant differences among all three cultivars (Fig 10, Tables Supplementary 8 in S1 text). Pinot Noir exhibited the highest estimated relative titre, with a fold change of 16.79 relative to Merlot (95% CI: 12.16-23.18), followed by Chasselas (5.38-fold relative to Merlot, 95% CI: 3.63-7.97). Merlot was used as the reference cultivar (1.00, 95% CI: 0.71-1.44).

**Fig 10.**
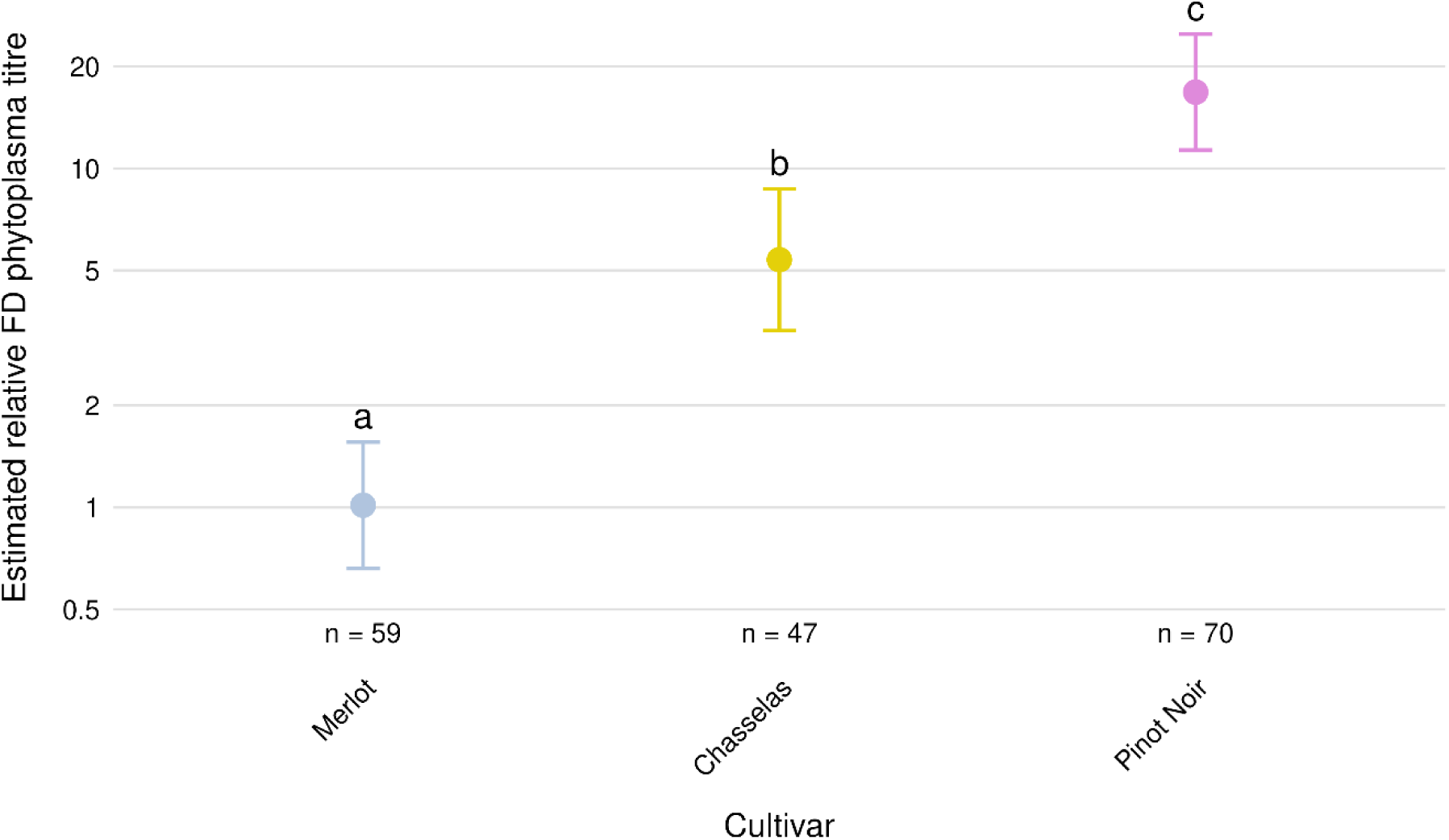
Model-estimated FDp titre expressed as fold change relative to Merlot in field samples of Chasselas and Pinot Noir. Points represent estimated marginal means obtained from a Gaussian model fitted on log-transformed data, and error bars indicate 95% confidence intervals. Fold changes were calculated using the ΔΔCt method, with Merlot used as the reference cultivar. Different letters indicate significant differences among cultivars based on Tukey-adjusted pairwise comparisons (α = 0.05).

## 4. Discussion

Because FD is a vector-borne disease, its incidence under vineyard conditions reflects not only the plant’s susceptibility to phytoplasma infection, but also plant-vector interactions and local epidemiological conditions. Consequently, assessing cultivar susceptibility requires complementary laboratory and field approaches that capture these different components.

In this study, we combined controlled transmission experiments with multi-scale field analyses to assess the susceptibility of the main Swiss grapevine cultivars to FD. The laboratory methodology differed slightly from previous studies (29,30), as we used cuttings instead of *ex vitro* plants to better approximate field conditions, given that *ex vitro Vitis vinifera* plants may retain juvenile traits (38). Nevertheless, in our experiment, the positions of Merlot and Cabernet Sauvignon at opposite ends of the susceptibility spectrum, as well as those of Chardonnay and Pinot Noir, were consistent with previously reported susceptibility patterns (29), supporting the relevance of the experimental approach.

The field study was based on up to ten years of surveillance data, covering more than 3,000 ha of vineyards divided into more than 49,000 plots and including more than 8,000 FD+ cases. To our knowledge, this is the first large-scale field analysis specifically addressing cultivar susceptibility to FD using complementary spatial scales, an approach that is particularly relevant in plant disease epidemiology because observed spatial patterns and their interpretation can depend on the scale of analysis (39,40). Among cultivars represented at both the macro and meso scales, Gamay showed a change in relative ranking. This result should be interpreted cautiously, as the meso-scale dataset contained substantially fewer hectares and FD+ cases for Gamay, as well as for Merlot and Gamaret, resulting in greater uncertainty around the estimated incidences. In contrast, the higher FD incidence in Pinot Noir than in Chasselas persisted across progressively finer spatial scales, including the micro scale, indicating that this cultivar effect was not solely driven by spatial variation in disease pressure or host distribution. The case study further supports this interpretation in a particularly clear epidemiological setting: despite exposure to the same identified inoculum source in a neighbouring Cabernet Dorsa vineyard, FD incidence was much higher in Pinot Noir than in Chasselas.

For most cultivars, laboratory and field assessments showed broadly consistent susceptibility patterns. The main exceptions were the two predominant Swiss cultivars, Pinot Noir and Chasselas. Field analyses consistently showed a higher FD incidence in Pinot Noir than in Chasselas, whereas no difference in infection probability was detected under controlled inoculation after accounting for the number of infectious *S. titanus*. Therefore, under the inoculation conditions used in this study, FDp established with similar probability in both cultivars. A similar discrepancy between the field and the laboratory was observed by Ripamonti *et al.* (30). They detected no significant difference in infection incidence between Barbera and Nebbiolo under laboratory conditions, whereas Roggia *et al.* (22) found that Nebbiolo was less susceptible than Barbera in the field. On the other hand, Eveillard *et al.* (29) had consistent results between both environments for Merlot and Cabernet Sauvignon in Bordeaux. Together, these studies highlight the value of combining controlled inoculation experiments with field epidemiological analyses, as each approach captures different components of cultivar susceptibility. Reliance on either approach alone may therefore provide an incomplete assessment.

The discrepancy observed between field and controlled inoculation conditions was also reflected in the analysis of relative phytoplasma titres: field samples from Pinot Noir exhibited higher relative titres than those from Chasselas (Fig 10), whereas no difference in this parameter was detected under controlled conditions (Fig 3). Again, these results indicate that, under the same inoculation pressure, FDp accumulated to similar relative titres in Chasselas and Pinot Noir. This contrasts with the observations of Eveillard et al. (29), who observed different phytoplasma titres between Merlot and Cabernet Sauvignon, both in the field and in the laboratory, consistent with more limited phytoplasma multiplication in Merlot.

In addition, the mortality of *S. titanus* during the one-week inoculation period differed significantly on the two cultivars. The estimated mortality probability of *S. titanus* was 16.8 percentage points lower on Pinot Noir than on Chasselas, and this difference was statistically significant. Our findings are consistent with those of Eveillard *et al.* (29) and Ripamonti *et al.* (30), who also reported cultivar-dependent differences in *S. titanus* mortality, with Merlot exhibiting higher mortality than the other cultivars (although only for grafted plants in the latter study). Furthermore, Ripamonti *et al.* (11) showed that *S. titanus* fitness differs among grapevine cultivars and broadly corresponds to cultivar-specific FD susceptibility patterns reported for the Piedmont region.

Taken together, our results indicate that, under the same inoculum pressure, FDp established with similar probability and accumulated to similar relative titres in Chasselas and Pinot Noir. Moreover, no major differences were detected in the distribution of the phytoplasma within the plant under controlled conditions, providing no evidence for the compartmentalisation phenomenon described by Casarin *et al.* in Tocai Friulano (18), and by Eveillard *et al*. (29) and ourselves in Merlot. These findings indicate that the contrasting field susceptibility of Chasselas and Pinot Noir is more likely to arise from differences in plant-vector interactions than from differences in host susceptibility to phytoplasma infection.

The higher mortality of *S. titanus* observed on Chasselas suggests that this cultivar may be a less suitable host for the vector than Pinot Noir, although this interpretation should be confirmed in experiments specifically designed to quantify cultivar effects on vector performance. Nevertheless, the reduced survival observed on Chasselas is consistent with the hypothesis that this cultivar expresses a higher level of resistance to *S. titanus*. If this effect also occurs under vineyard conditions, reduced vector survival could contribute to lower vector abundance and, consequently, lower FD transmission pressure in Chasselas vineyards. The mechanisms underlying this putative resistance remain unknown and may involve differences in host acceptance, feeding behaviour, nutritional suitability or plant defence responses. Further studies combining field surveys with behavioural, feeding and performance assays are needed to determine how cultivar traits affect *S. titanus* and whether these effects contribute to cultivar-dependent FD incidence.

In conclusion, our findings illustrate that controlled inoculation experiments alone may not fully predict cultivar susceptibility under vineyard conditions, highlighting the importance of integrating laboratory and field approaches when assessing susceptibility to vector-borne plant diseases. Although Chasselas and Pinot Noir exhibited similar susceptibility to FDp following controlled inoculation, Pinot Noir consistently showed substantially higher disease incidence under vineyard conditions. Together with the higher mortality of *S. titanus* observed on Chasselas during the inoculation period, these findings support the hypothesis that cultivar-specific differences in host suitability for the insect vector, rather than intrinsic susceptibility to phytoplasma infection, contribute to field disease incidence. If this hypothesis is confirmed and the mechanisms underlying cultivar-specific differences in host suitability for *S. titanus* are elucidated, the plant traits responsible for these differences could represent useful targets for grapevine breeding programmes aimed at reducing vector suitability and contributing to more sustainable management of flavescence dorée.

## Data availability

The datasets underlying the laboratory experiments and field analyses are publicly available on Zenodo (DOI: 10.5281/zenodo.21885421). The analysis code used to reproduce the statistical analyses and generate the tables and figures reported in the manuscript is publicly available on Zenodo (DOI: 10.5281/zenodo.21930033).

The original georeferenced vineyard surveillance data underlying Figure 1 and S2 Fig, S3 Fig and S4 Fig cannot be made publicly available due to confidentiality restrictions associated with georeferenced phytosanitary surveillance data. These data were provided by the cantonal phytosanitary services and are not owned by the authors. Access to the original data may be requested directly from the relevant cantonal phytosanitary services, subject to their applicable data-access requirements.

## Acknowledgments

The authors sincerely thank Larisa Grosu-Duchêne, Marc Passerat, Éric Remolif, and Stefan Kellenberger (Agroscope, Changins) for producing and providing the grapevine and broad bean plants used in the experiments. We are also grateful to Laure Apothéloz-Perret-Gentil (Agroscope, Changins) for providing data used in the field analysis of FDp relative titre, to Victor Rueda Ayala (Agroscope, Changins) for his valuable statistical advice, to Matthieu Wilhelm (Agroscope, Changins) for his guidance on statistical reporting and the preparation of reproducible Quarto documents, and to Yvonne Fuchs (Agroscope, Bern) for her guidance on open research data publication. We further thank Domenico Bosco and his team at the University of Turin and CNR-IPSP, Turin, for providing the initial colony of *Euscelidius variegatus* used to establish our laboratory colony.

## Notes

### Competing Interest Statement

The authors have declared no competing interest.

### Summary of Updates

Submission of supplementary text and figures

