## Supplementary material for "Integrated field and laboratory assessment of Swiss grapevine cultivar susceptibility to flavescence dorée reveals a central role for plant-vector interactions": S1 text

**Table Supplementary 1. Overview of the grapevine transmission assays conducted under controlled conditions in 2023 and 2025 and retained for susceptibility probability analysis**. For each cultivar and experimental replicate, the table reports the year, experimental session (inoculation block in which a subset of cultivars was exposed simultaneously to the same batch of infectious insects), and the percentage of infectious insects.

| Cultivar | Year | Experimental replicate | Experimental session | Infectious insects (%) |
|---|---|---|---|---|
| **Chasselas** | 2023 | 1 | S02 | 75 |
| **Chasselas** | 2023 | 2 | S03 | 75 |
| **Chasselas** | 2023 | 3 | S06 | 92.5 |
| **Chasselas** | 2025 | 4 | S09 | 92.9 |
| **Chasselas** | 2025 | 5 | S10 | 80 |
| **Chardonnay** | 2023 | 3 | S05 | 88.5 |
| **Chardonnay** | 2025 | 4 | S07 | 92.9 |
| **Chardonnay** | 2025 | 5 | S09 | 87.5 |
| **Cabernet Sauvignon** | 2025 | 1 | S07 | 92.9 |
| **Cabernet Sauvignon** | 2025 | 2 | S08 | 92.6 |
| **Cabernet Sauvignon** | 2025 | 3 | S09 | 95 |
| **Cabernet Sauvignon** | 2025 | 4 | S10 | 78.6 |
| **Gamaret** | 2023 | 1 | S03 | 95 |
| **Gamaret** | 2023 | 2 | S05 | 100 |
| **Gamaret** | 2025 | 3 | S07 | 89.3 |
| **Gamaret** | 2025 | 4 | S08 | 87.5 |
| **Gamay** | 2023 | 1 | S03 | 85.2 |
| **Gamay** | 2023 | 2 | S05 | 100 |
| **Gamay** | 2025 | 3 | S07 | 96.4 |
| **Gamay** | 2025 | 4 | S08 | 88.4 |
| **Merlot** | 2023 | 2 | S04 | 77.8 |
| **Merlot** | 2025 | 3 | S07 | 96.4 |
| **Merlot** | 2025 | 4 | S09 | 87.2 |
| **Pinot Noir** | 2023 | 1 | S02 | 79.2 |
| **Pinot Noir** | 2023 | 2 | S03 | 85.2 |
| **Pinot Noir** | 2023 | 3 | S06 | 85 |
| **Pinot Noir** | 2025 | 4 | S09 | 88.9 |
| **Pinot Noir** | 2025 | 5 | S10 | 96.2 |

**Table Supplementary 2 Results of the laboratory analysis of grapevine susceptibility to flavescence dorée following controlled insect-mediated inoculation.**

**Table Sup2A Parameter estimates of the final complementary log-log binomial model used to analyse the probability of plant infection following controlled insect-mediated inoculation**. Merlot was used as the reference cultivar. Estimates are reported on the complementary log-log scale.

| Parameter | Estimate | SE | z | P |
|---|---|---|---|---|
| **Intercept (Merlot)** | -2.5510 | 0.4098 | -6.225 | <0.001 |
| **Chasselas** | 0.6776 | 0.4775 | 1.419 | 0.156 |
| **Chardonnay** | 1.1454 | 0.4937 | 2.320 | 0.020 |
| **Cabernet Sauvignon** | 1.3383 | 0.4725 | 2.832 | 0.005 |
| **Gamaret** | 1.0552 | 0.4801 | 2.198 | 0.028 |
| **Gamay** | 0.9692 | 0.4791 | 2.023 | 0.043 |
| **Pinot Noir** | 0.6719 | 0.4814 | 1.396 | 0.163 |

**Table Sup2B Model-based infection probabilities (%) for each cultivar at a standardized exposure of four infected insects per plant.** Values are presented with 95% confidence intervals (95% CI). P-values correspond to planned contrasts between each cultivar and the reference cultivar Merlot, based on estimated marginal means and adjusted using the Dunnett's method.

| Cultivar | Infected / exposed plants | Infection probability (%) | 95% CI | Delta P vs Merlot (percentage points) | Dunnett adjusted p-value vs Merlot |
|---|---|---|---|---|---|
| **Merlot** | 6 / 26 | 26.8 | 13.04-50.17 | Ref. | Ref. |
| **Pinot Noir** | 16 / 40 | 45.71 | 31.08-63.30 | 18.91 | 0.457 |
| **Chasselas** | 17 / 43 | 45.9 | 31.62-62.96 | 19.1 | 0.436 |
| **Gamay** | 17 / 34 | 56.06 | 39.69-73.76 | 29.26 | 0.113 |
| **Gamaret** | 17 / 31 | 59.19 | 42.25-76.85 | 32.39 | 0.066 |
| **Chardonnay** | 14 / 25 | 62.5 | 43.55-81.41 | 35.7 | 0.048 |
| **Cabernet Sauvignon** | 20 / 31 | 69.56 | 52.77-84.83 | 42.76 | 0.004 |

**Table Supplementary 3 Results of the laboratory analysis of relative FD phytoplasma titre among grapevine cultivars following controlled insect-mediated inoculation.**

**Table Sup3A Parameter estimates of the final Gaussian model used to analyse relative FD phytoplasma titre following controlled insect-mediated inoculation.** Merlot was used as the reference cultivar. Estimates are reported on the log-transformed response scale.

| Parameter | Estimate | SE | z | P |
|---|---|---|---|---|
| **Intercept (Merlot)** | -1.338 | 1.298 | -1.031 | 0.302 |
| **Chasselas** | -0.976 | 1.510 | -0.646 | 0.518 |
| **Chardonnay** | 1.794 | 1.551 | 1.157 | 0.247 |
| **Cabernet Sauvignon** | 0.741 | 1.480 | 0.501 | 0.616 |
| **Gamaret** | 1.037 | 1.510 | 0.687 | 0.492 |
| **Gamay** | 2.000 | 1.510 | 1.325 | 0.185 |
| **Pinot Noir** | 0.151 | 1.522 | 0.100 | 0.921 |

**Table Sup3B Estimated relative FD phytoplasma titre in grapevine cultivars following controlled insect-mediated inoculation.** Values are estimated marginal means expressed as ΔΔCt-derived fold changes relative to the reference cultivar Merlot, with 95% confidence intervals (CI). Statistical groups are based on Dunnett-adjusted comparisons against the reference cultivar (Merlot). Cultivars sharing the same letter did not differ significantly from Merlot ($P > 0.05$).

| Cultivar | Estimated fold change (95% CI) | Statistical group |
|---|---|---|
| **Merlot (reference)** | 1.05 (0.08-13.78) | a |
| **Chasselas** | 0.40 (0.09-1.83) | a |
| **Chardonnay** | 6.31 (1.17-34.06) | a |
| **Cabernet Sauvignon** | 2.20 (0.54-9.02) | a |
| **Gamaret** | 2.96 (0.64-13.66) | a |
| **Gamay** | 7.75 (1.68-35.79) | a |
| **Pinot Noir** | 1.22 (0.25-5.91) | a |

**Table Supplementary 4 Results of the laboratory analysis of insect mortality across grapevine cultivars.**

**Table Sup4A Parameter estimates of the final binomial model used to analyse insect mortality.** Chasselas, control treatment and session S02 were used as the reference levels. Estimates are reported on the logit scale.

| Parameter | Estimate | SE | z | P |
|---|---|---|---|---|
| **Intercept (Chasselas, control, session S02)** | -1.354 | 0.401 | -3.377 | <0.001 |
| **Chardonnay** | -0.621 | 0.442 | -1.405 | 0.16 |
| **Cabernet Sauvignon** | -0.865 | 0.387 | -2.234 | 0.025 |
| **Gamaret** | 0.02 | 0.427 | 0.047 | 0.963 |
| **Gamay** | -0.527 | 0.413 | -1.275 | 0.202 |
| **Merlot** | 0.853 | 0.525 | 1.625 | 0.104 |
| **Pinot Noir** | -0.679 | 0.299 | -2.267 | 0.023 |
| **Inoculated treatment** | 0.854 | 0.229 | 3.733 | <0.001 |
| **Session S03** | 0.994 | 0.447 | 2.225 | 0.026 |
| **Session S04** | 0.109 | 0.752 | 0.145 | 0.885 |
| **Session S05** | 0.698 | 0.543 | 1.285 | 0.199 |
| **Session S06** | 1.731 | 0.455 | 3.807 | <0.001 |
| **Session S07** | 1.108 | 0.512 | 2.164 | 0.03 |
| **Session S08** | 1.91 | 0.538 | 3.552 | <0.001 |
| **Session S09** | 2.264 | 0.468 | 4.833 | <0.001 |
| **Session S10** | 1.998 | 0.455 | 4.389 | <0.001 |

**Table Sup4B Model-based mortality probabilities (%) of infectious insects for each cultivar averaged across treatment and experimental session.** Values are presented with 95% confidence intervals (CI). Absolute differences (ΔP) are expressed in percentage points relative to the reference cultivar Merlot. Fold change represents the ratio of mortality probabilities compared with Merlot. P-values correspond to Dunnett-adjusted planned contrasts versus Merlot.

| Cultivar | Mortality probability (%) (95% CI) | Delta probability vs Merlot (percentage points) | Dunnett-adjusted p-value vs Merlot |
|---|---|---|---|
| **Merlot** | 75.5 (58.6-87.1) | - | - |
| **Pinot Noir** | 40.0 (28.5-52.8) | -35.5 | 0.017 |
| **Chasselas** | 56.8 (43.8-69.0) | -18.7 | 0.384 |
| **Gamay** | 43.7 (30.5-57.9) | -31.8 | 0.034 |
| **Gamaret** | 57.3 (42.4-71.0) | -18.2 | 0.385 |
| **Chardonnay** | 41.4 (27.0-57.4) | -34.1 | 0.018 |
| **Cabernet Sauvignon** | 35.7 (24.2-49.0) | -39.9 | 0.003 |

**Table Supplementary 5 Results of the macro-scale analysis of flavescence dorée incidence across the main grapevine cultivars.**

**Table Sup5A Parameter estimates of the final NB1 negative binomial model used to analyse the density of FD-positive grapevines in the macro-scale analysis.** Chasselas was used as the reference cultivar. Estimates are reported on the log link scale.

| Parameter | Estimate | SE | z | P |
|---|---|---|---|---|
| **Intercept (Chasselas)** | -1.316 | 0.103 | -12.814 | <0.001 |
| **Pinot Noir** | 2.255 | 0.142 | 15.886 | <0.001 |
| **Gamay** | 2.981 | 0.167 | 17.803 | <0.001 |
| **Sylvaner** | -2.176 | 0.477 | -4.563 | <0.001 |
| **Arvine** | -0.614 | 0.358 | -1.713 | 0.087 |

**Table Sup5B Observed and model-based FD-positive grapevine density among the five main grapevine cultivars included in the macro-scale study.** Observed values correspond to vineyard area, the total number of FD-positive grapevines, and the observed FD+ density (FD+ plants $ha^{-1}$). Estimated values are expressed as FD-positive grapevines per hectare with 95% confidence intervals (95% CI). Different letters indicate significant differences based on Tukey-adjusted pairwise comparisons (α = 0.05).

| Cultivar | Surface (ha) | FD+ | Observed FD+ ha-1 | Estimated FD+ ha-1 (95% CI) | Statistical group |
|---|---|---|---|---|---|
| **Sylvaner** | 205 | 7 | 0.03 | 0.03 (95% CI: 0.01-0.08) | a |
| **Arvine** | 171 | 15 | 0.09 | 0.15 (95% CI: 0.07-0.28) | ab |
| **Chasselas** | 1452 | 518 | 0.36 | 0.27 (95% CI: 0.22-0.33) | b |
| **Pinot Noir** | 1000 | 4175 | 4.18 | 2.56 (95% CI: 2.11-3.10) | c |
| **Gamay** | 375 | 3408 | 9.08 | 5.29 (95% CI: 4.08-6.85) | d |

**Table Supplementary 6 Results of the meso-scale analysis of flavescence dorée incidence among the five main grapevine cultivars included in the 500 m buffer zone study in the canton of Vaud.**

**Table Sup6A Parameter estimates of the final NB2 negative binomial model used to analyse FD-positive grapevine density in the meso-scale study.** Merlot was used as the reference cultivar. Estimates are reported on the log link scale.

| Term | Estimate | SE | z | P value |
|---|---|---|---|---|
| **Intercept (Merlot)** | 0.221 | 0.530 | 0.416 | 0.677 |
| **Pinot Noir** | 2.943 | 0.573 | 5.135 | <0.001 |
| **Chasselas** | -0.088 | 0.543 | -0.161 | 0.872 |
| **Gamay** | 2.271 | 0.614 | 3.700 | <0.001 |
| **Gamaret** | 2.615 | 0.653 | 4.007 | <0.001 |

**Table Sup6B Observed and model-based FD-positive grapevine density among the five main grapevine cultivars included in the meso-scale study (500 m buffer zones, canton of Vaud).** Observed values correspond to at-risk vineyard surface (ha), the total number of FD-positive grapevines, and the observed FD+ density (FD+ plants $ha^{-1}$). Estimated values are expressed as FD-positive grapevines per hectare with 95% confidence intervals (95% CI). Different letters indicate significant differences based on Tukey-adjusted pairwise comparisons (α = 0.05).

| Cultivar | At-risk surface (ha) | FD+ | Observed FD+ $ha^{-1}$ | Estimated FD+ $ha^{-1}$ (95% CI) | Statistical group |
|---|---|---|---|---|---|
| **Chasselas** | 528 | 493 | 0.93 | 1.14 (95% CI: 0.91-1.44) | a |
| **Merlot** | 13 | 12 | 0.92 | 1.25 (95% CI: 0.44-3.53) | a |
| **Gamay** | 31 | 291 | 9.39 | 12.08 (95% CI: 6.60-22.14) | b |
| **Gamaret** | 17 | 402 | 23.65 | 17.04 (95% CI: 8.09-35.90) | b |
| **Pinot Noir** | 75 | 1,739 | 23.19 | 23.66 (95% CI: 15.47-36.20) | b |

**Table Supplementary 7 Results of the micro-scale analysis of flavescence dorée incidence in adjacent Pinot Noir and Chasselas vineyards in the canton of Vaud.**

**Table Sup7A Parameter estimates of the final NB2 negative binomial mixed model used to analyse FD-positive grapevine density in the micro-scale study.** Chasselas was used as the reference cultivar. Estimates are reported on the log link scale.

| Term | Estimate | SE | z | P value |
|---|---|---|---|---|
| **Intercept (Chasselas)** | 1.459 | 0.328 | 4.448 | <0.001 |
| **Pinot Noir** | 1.882 | 0.332 | 5.669 | <0.001 |

**Table Sup7B Observed and model-based FD-positive grapevine density among the two main grapevine cultivars included in the micro-scale study (15 m buffer zones along Pinot Noir-Chasselas boundaries, canton of Vaud).** Observed values correspond to vineyard surface, the total number of FD-positive grapevines, and the observed FD+ density (FD+ plants $ha^{-1}$). Estimated values are expressed as FD-positive grapevines per hectare with 95% confidence intervals (95% CI). Different letters indicate significant differences among cultivars.

| Cultivar | Surface (ha) | FD+ | Observed FD+ $ha^{-1}$ | Estimated FD+ $ha^{-1}$ (95% CI) | Statistical group |
|---|---|---|---|---|---|
| **Chasselas** | 15.70 | 146 | 9.30 | 4.30 (95% CI: 2.26-8.19) | a |
| **Pinot Noir** | 9.46 | 738 | 78.02 | 28.27 (95% CI: 15.86-50.39) | b |

**Table Supplementary 8 Results of the relative FD phytoplasma titre in field samples of Merlot, Chasselas and Pinot Noir.** Relative FD phytoplasma titre was expressed as fold change relative to Merlot using the ΔΔCt method.

**Table Sup8A Parameter estimates of the final log-Gaussian model.** Merlot was used as the reference cultivar. Estimates are reported on the log-transformed response scale.

| Term | Estimate | SE | z | P value |
|---|---|---|---|---|
| **Intercept (Merlot)** | 0.015 | 0.178 | 0.083 | 0.934 |
| **Chasselas** | 1.667 | 0.267 | 6.240 | <0.001 |
| **Pinot Noir** | 2.806 | 0.242 | 11.617 | <0.001 |

**Table Sup8B Estimated marginal means back-transformed to the response scale and expressed as ΔΔCt-derived fold change relative to Merlot with 95% confidence intervals (95% CI).** Different letters indicate significant differences among cultivars based on Tukey-adjusted pairwise comparisons.

| Cultivar | Estimated fold change (95% CI) | Statistical group |
|---|---|---|
| **Merlot (reference)** | 1.01 (0.71-1.44) | a |
| **Chasselas** | 5.38 (3.63-7.97) | b |
| **Pinot Noir** | 16.79 (12.16-23.18) | c |
