## Supplementary material for "Integrated field and laboratory assessment of Swiss grapevine cultivar susceptibility to flavescence dorée reveals a central role for plant-vector interactions": suppl_figures

**Supplementary Figure 1** Sampling scheme used for the transmission assays. Sixteen weeks after the start of the transmission assays, four leaves were collected from each plant whenever possible: the inoculated leaf (IL), one leaf between the 3rd and 5th upper leaves (proximal upper leaf, PUL), one leaf between the 8th and 10th upper leaves (distal upper leaf, DUL), and one leaf between the 3rd and 5th lower leaves (basal leaf, BL).

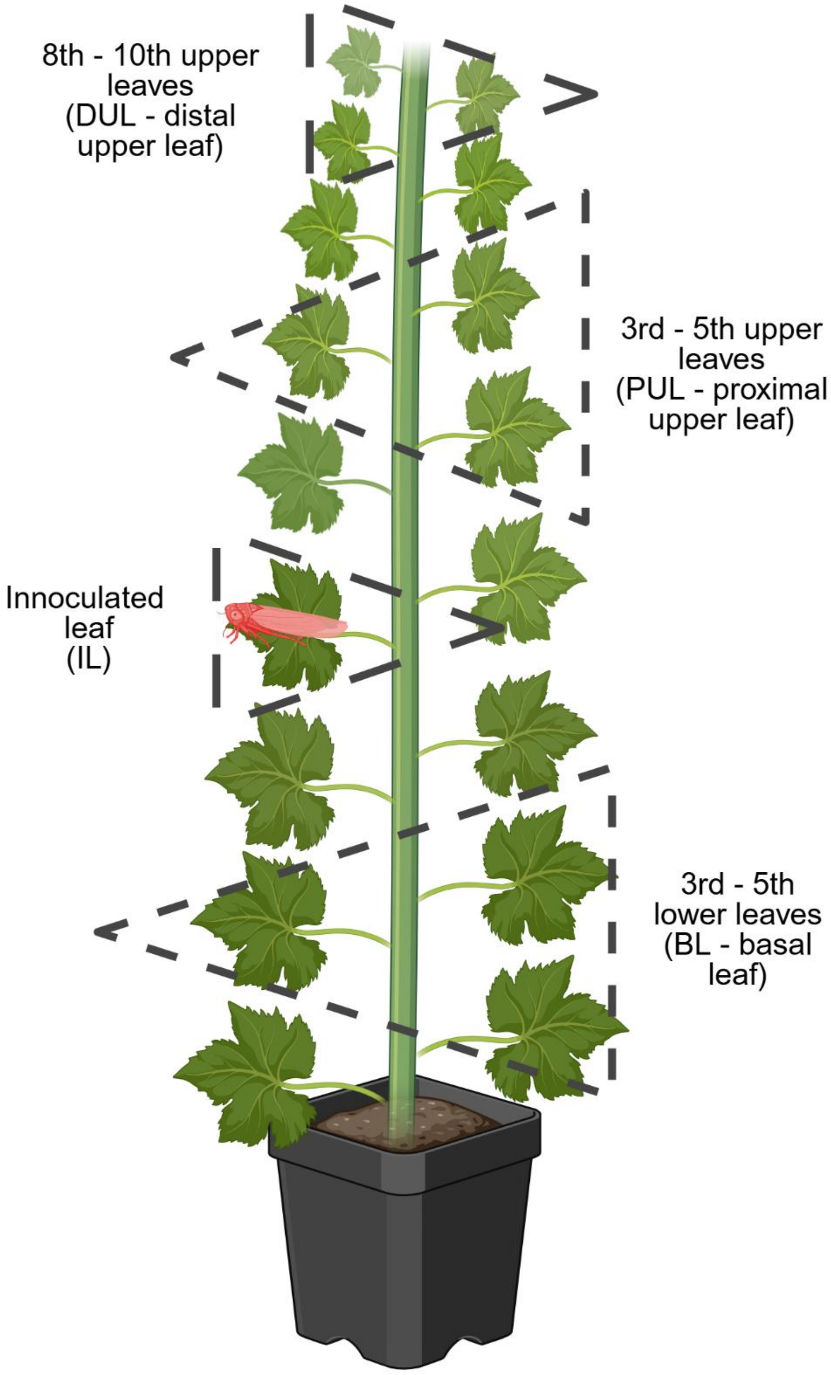

**Supplementary Figure 2** Vineyard areas in the cantons of Geneva, Vaud and Valais, where flavescence dorée is present. Vineyards included in the macro-scale analyses (Vaud and Valais) are shown in dark purple, whereas vineyard areas excluded from the analyses are shown in light purple.

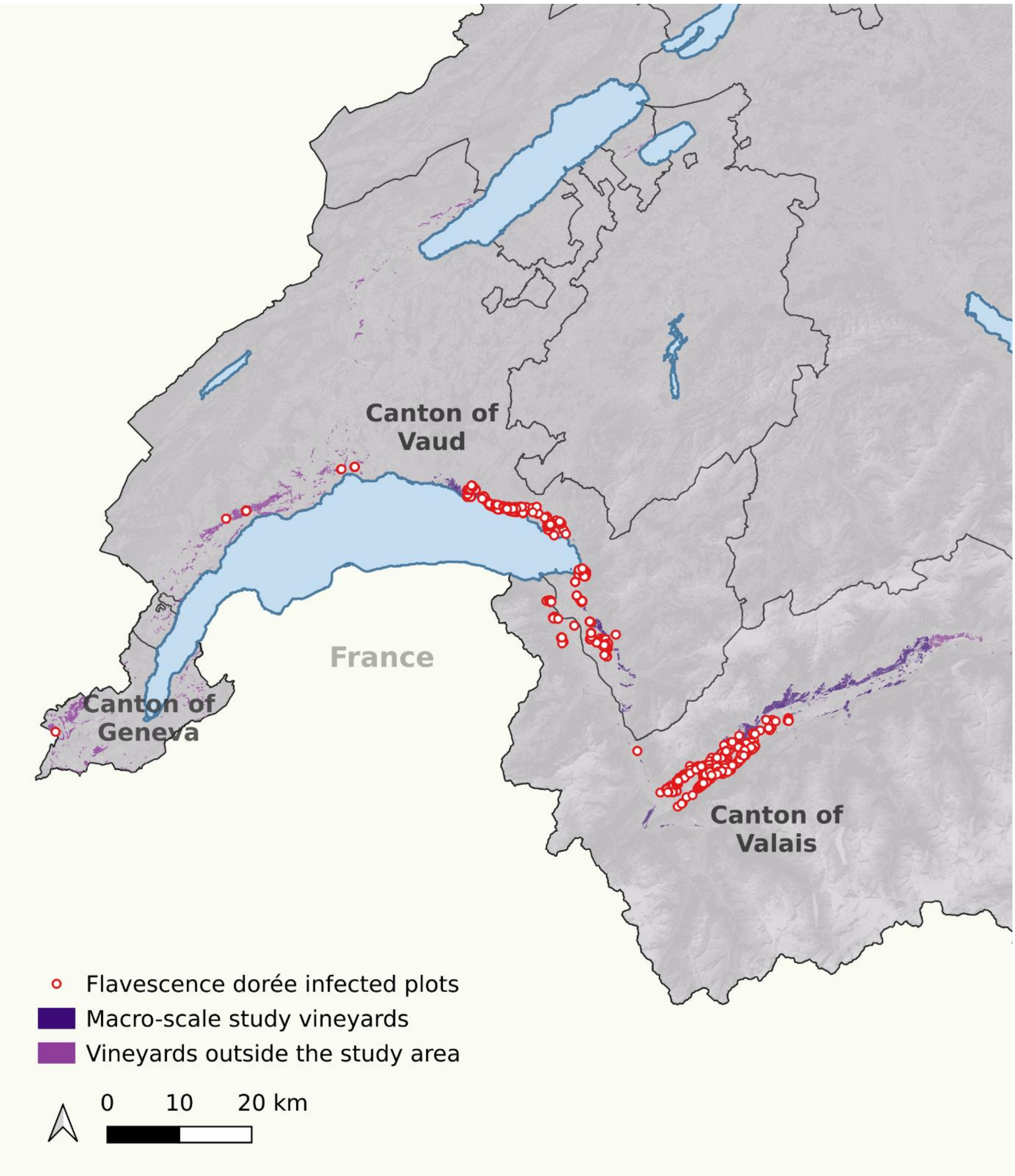

**Supplementary Figure 3** Representative example of the meso-scale analysis, showing the 500 m buffers generated around FD-positive grapevines. Only vineyard areas located within the 500 m buffers were included in the analyses.

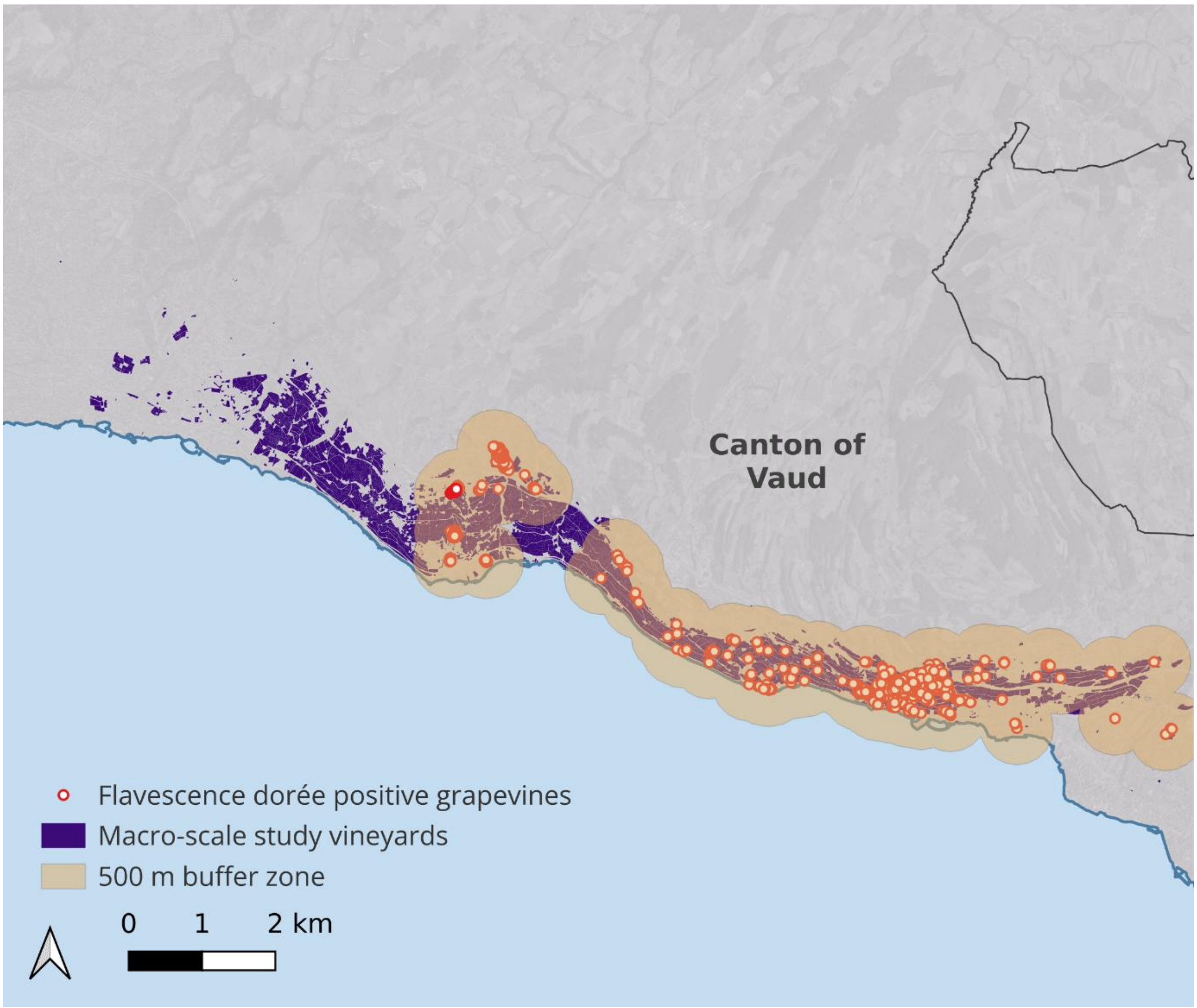

**Supplementary Figure 4** Representative example of the micro-scale analysis, showing the 15 m buffers generated on both sides of contact zones between adjacent Chasselas and Pinot Noir vineyards. Only 15 m buffers containing at least one FD-positive grapevine were included in the analyses.

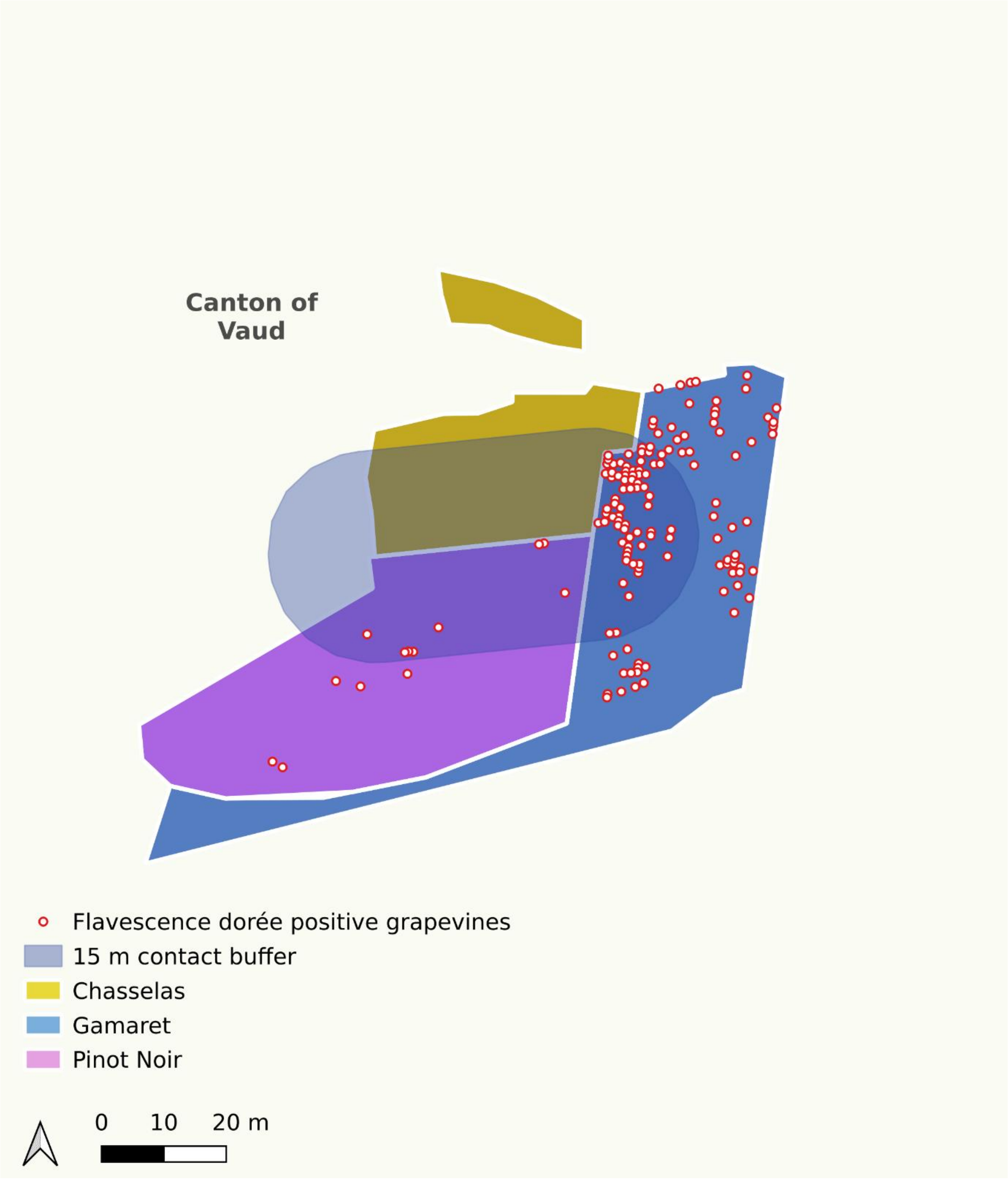
